# ERH regulates type II interferon immune signaling through post-transcriptional regulation of *JAK2* mRNA

**DOI:** 10.1101/2024.08.20.607899

**Authors:** Adrian Soderholm, Milica Vunjak, Melanie De Almeida, Niko Popitsch, Nadezda Podvalnaya, Pablo Araguas-Rodriguez, Sara Scinicariello, Emily Nischwitz, Falk Butter, René Ketting, Stefan L. Ameres, Michaela Müller-McNicoll, Johannes Zuber, Gijs A. Versteeg

## Abstract

Type II interferon (IFNγ) signaling is essential for innate immunity and critical for effective immunological checkpoint blockade in cancer immunotherapy. Genetic screen identification of post-transcriptional regulators of this pathway has been challenging since such factors are often essential for cell viability. Here, we utilize our inducible CRISPR/Cas9 approach to screen for key post-transcriptional regulators of IFNγ signaling, and in this way identify ERH and the ERH-associated splicing and RNA export factors MAGOH, SRSF1, and ALYREF. Loss of these factors impairs post-transcriptional mRNA maturation of *JAK2*, a crucial kinase for IFNγ signaling, resulting in abrogated JAK2 protein levels and diminished IFNγ signaling. Further analysis highlights a critical role for ERH in preventing intron retention in AU-rich regions in specific transcripts, such as *JAK2*. This regulation is markedly different from previously described retention of GC-rich introns. Overall, these findings reveal that post-transcriptional *JAK2* processing is a critical rate-limiting step for the IFNγ-driven innate immune response.

## Introduction

Interferons (IFNs) are critical cytokines in the innate immune system, mediating the first line of defense against diverse pathogens (Castro *et al*, 2018; Philips *et al*, 2022). These cytokines fall into three main categories: type I (IFNα and IFNβ), type II (IFNγ), and type III (IFNλ). Upon binding specific cell surface receptors, IFNs trigger signaling cascades that activate Janus kinase (JAK) and signal transducer and activator of transcription (STAT) pathways (Shuai *et al*, 1992, 1993; Decker *et al*, 1991; Pellegrini *et al*, 1989; Krolewski *et al*, 1990; Harpur *et al*, 1992; Wilks *et al*, 1991; Firmbach-Kraft *et al*, 1990). This activation results in the transcription of hundreds of interferon-stimulated genes (ISGs) that encode proteins with antiviral, antiproliferative, and immunomodulatory properties (Levy *et al*, 1986; Schoggins *et al*, 2011).

Similar to type I and type III IFNs, IFNγ signaling can drive antiviral gene expression but is primarily involved in anti-bacterial responses (Ivashkiv, 2018). IFNγ signaling is critically dependent on the JAK/STAT pathway composed of two receptor heterodimers (IFNGR1 and IFNGR2) and activation of the associated kinases (JAK1 and JAK2) that phosphorylate the transcription factor STAT1, which in turn drives ISG expression (Fig. 1A). In this way, IFNγ activates macrophages, thus promoting phagocytosis and the production of reactive oxygen and nitrogen species – critical components for eliminating intracellular bacterial pathogens (Ivashkiv, 2018).

**Figure 1.**
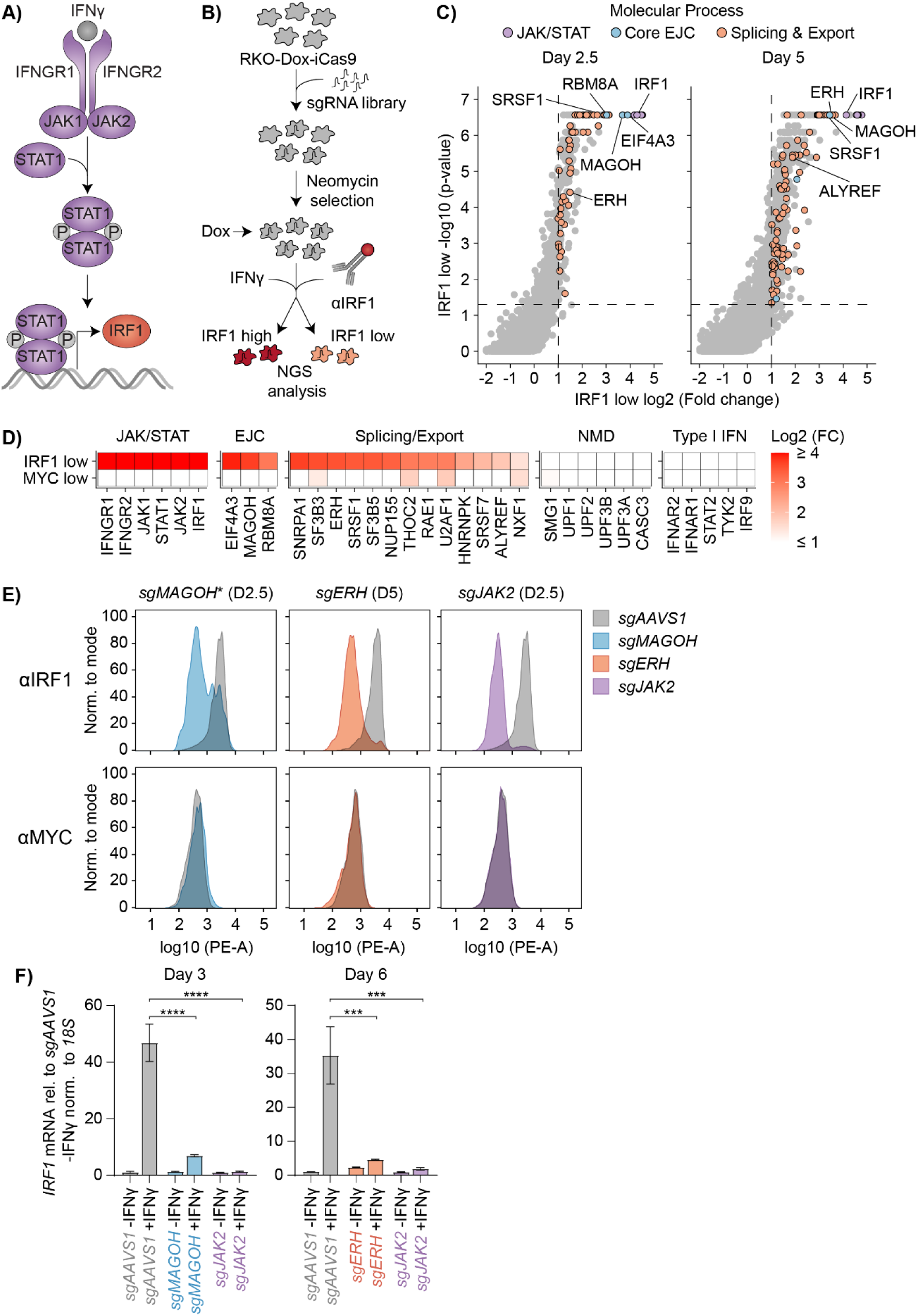
Identification of ERH and other novel positive regulators of IFNγ signaling by genome-wide genetic screening. **(A)** Schematic of the IFNγ-induced JAK/STAT signaling pathway, which stimulates expression of many genes, including IRF1. (**B**) Overview of FACS-based CRISPR-Cas9 knockout screen. Human RKO cells with dox-inducible iCas9 were transduced with a lentiviral genome-wide *sgRNA* library. Cas9 expression was induced for 2.5 or 5 days, after which cells were treated with IFNγ, and IRF1 induction detected by intracellular staining. Cells with the lowest or highest IRF1 levels were collected by FACS, and disrupted genes were identified by analyzing sgRNA-targeted coding sequences. (**C**) sgRNA enrichment in the IRF1^low^ cell population was plotted. Dashed lines indicate significance (p ≤ 0.05) and enrichment (log2 fold change ≥ 1). Significantly enriched genes involved in the JAK/STAT pathway, the exon junction complex, or RNA splicing and export are highlighted. (**D**) Heatmap of selected IRF1 regulators as in (C) or MYC regulators (de Almeida *et al*, 2021) involved in JAK/STAT signaling, the EJC, splicing and export, nonsense mediated decay, or type I interferon signaling. For each gene, the time point with the strongest enrichment is plotted. (**E**) RKO-iCas9 cells were transduced with vectors expressing the indicated sgRNAs. After 2.5 or 5 days of dox-induced Cas9 expression, cells were stimulated with IFNγ, and endogenous IRF1 or MYC were detected by intra-cellular staining and flow cytometry. Representative sample from 2 (n = 2) or *1 (n = 1) biological replicates are shown. (**F**) In parallel, *IRF1* mRNA levels were measured by RT-qPCR. Data represent the mean and sd; n = 3 biological replicates. Two-tailed t-test with Benjamini-Hochberg correction (*p ≤ 0.05; **p ≤ 0.01; ***p ≤ 0.001; ****p ≤ 0.0001).

Moreover, IFNγ signaling is known to be key for effective checkpoint inhibition that is used in cancer immunotherapy to relieve immune cells of negative signals from check point proteins such as CTLA4 and PD1. In this context, non-responders to anti-CTLA4 therapy were found to have defects in IFNγ signaling within their tumors (Gao *et al*, 2016). In addition, failure to respond to IFNγ and anti-PD1 melanoma therapy was associated with loss-of-function mutations in *JAK1* and *JAK2* in tumor biopsy samples (Zaretsky *et al*, 2016). Despite their numerous host-beneficial immunomodulatory functions, JAK1/JAK2 activity resulting from IFNγ signaling requires proper regulation. This is exemplified by the fact that mutations in exon 14 of *JAK2* (V617F) result in its constitutive activation and leads to myeloproliferative neoplasms (James *et al*, 2005; Baxter *et al*, 2005; Kralovics *et al*, 2005).

To prevent aberrant activation, innate immune signaling is well-recognized to be tightly controlled at the post-translational level. For example, the phosphorylation of receptor-activated kinases critically regulates the key transcription factors driving type I IFN transcription (IRF3 and IRF7), as well as STAT1-containing transcription factor complexes driving ISG expression in both type I and type II IFN signaling (Xue *et al*, 2023b).

However, evidence is emerging that key factors of these pathways, as well as individual ISGs, are also post-transcriptionally regulated (O’Connor *et al*, 2015; Green *et al*, 2020; Lefaudeux *et al*, 2022; Yabas *et al*, 2015). For example, alternative splicing and intron retention (IR) are critical mechanisms that regulate mRNAs encoding key immune factors that modulate inflammatory responses (O’Connor *et al*, 2015; Yabas *et al*, 2015), and control macrophage development and function (Green *et al*, 2020; Wagner *et al*, 2021). Furthermore, mRNA export rates have been shown to complement mRNA decay rates to properly regulate innate immune responses (Lefaudeux *et al*, 2022). Overall, these examples highlight components of the post-transcriptional RNA processing machinery as critical factors for innate immunity.

RNA processing factors comprise a significant proportion of genes that are important for cellular viability, likely since they regulate large groups of mRNAs, including those needed for fundamental biological processes (Wang *et al*, 2015; Kwon *et al*, 2021). Their essential nature has made it difficult to systematically screen for these regulators. Here we set out to identify novel factors controlling IFNγ signaling, with a focus on post-transcriptional regulators. Given that many of these genes are essential for cell viability, we hypothesized that previous gene trap-mediated genetic screening (Brockmann *et al*, 2017) may have overlooked critical post-transcriptional factors that function in IFNγ signaling. Therefore, we utilized our established inducible CRISPR/Cas9 platform (de Almeida *et al*, 2021), which allows identification of cell-essential genes.

Using this approach, we identified Enhancer of Rudimentary Homolog (ERH) and its interacting splicing and export factors as novel positive regulators of IFNγ-induced signaling. These factors act through post-transcriptional regulation of *JAK2*: in their absence, JAK2 protein was lost, thereby abrogating IFNγ signaling. Our data suggest that post-transcriptional regulation of *JAK2* is rate-limiting and disproportionately affected by the disruption of these RNA regulatory factors. Furthermore, ERH was critical for preventing intron retention in *JAK2* and multiple other mRNAs, including several involved in DNA replication and the DNA damage response. Our results specifically reveal that ERH is essential for efficient splicing in AU-rich regions, contrasting with previously described intron retention of GC-rich introns. Overall, these findings highlight the importance of post-transcriptional *JAK2* processing by ERH and interacting factors in mounting an effective type II IFN response.

## Results

### Identification of novel positive regulators of IFNγ signaling by genome-wide genetic screening

To identify novel regulatory factors of the type II interferon signaling pathway, we performed a genome-wide genetic screen using our previously established time-controlled CRISPR/Cas9 screening approach (de Almeida *et al*, 2021; Michlits *et al*, 2020). We used an optimized doxycycline (dox)-inducible Cas9 (iCas9) human RKO cell line (de Almeida *et al*, 2021), where functional editing by Cas9 only occurs in the presence of dox (Fig. EV1A). This allows for temporal control of gene knockouts and thus the possibility to assess the effects of disrupting cell-essential genes (de Almeida *et al*, 2021).

A genome-wide lentiviral sgRNA library (Michlits *et al*, 2020) was transduced into the RKO-dox-iCas9 line at low multiplicity of transduction to ensure one integration per cell. Cells with single-gene disruptions were treated with IFNγ, and endogenous protein levels of the ISG IRF1 were used as a proxy for IFNγ signaling (Fig. 1A). IFNγ-treated knockout cells with the lowest or highest IRF1 protein levels were then collected by fluorescence-activated cell sorting (FACS) for further analysis (Fig. 1B). Two timepoints (2.5 and 5 days after Cas9 induction by dox) were used to allow the identification of factors independent of their turnover rate and whether they were essential for viability. The relative abundances of sgRNAs in collected cells were analyzed by next-generation sequencing (NGS) and compared to day-matched mean-distribution cell pools representing pre-enrichment library sgRNA frequencies (Fig. 1B). This comprehensive assay thus measures the effects of transcriptional to post-translational regulation on IFNγ signaling.

To identify factors required for IFNγ signaling, and hence IRF1 induction, we analyzed enriched sgRNAs in the IRF1^low^ cell population. As expected, IRF1 itself and all well-known JAK/STAT factors involved in propagating IFNγ-signaling (Fig. 1A) scored as top hits (Fig. 1C,D and Fig. EV1B), attesting to the validity of the screen. In contrast, factors which are critical for type I IFN signaling, but not for IFNγ-dependent responses, were not enriched (Fig. 1D), demonstrating its specificity.

To discriminate specific from more general effects, we compared our hits with those of a similar screen that successfully identified regulators of the transcription factor MYC (de Almeida *et al*, 2021). We reasoned that MYC would be a good control to filter out general RNA- and protein regulatory factors, since *MYC* mRNA and protein are rapidly turned over in cells (Lemm & Ross, 2002; Gregory & Hann, 2000; Ramsay *et al*, 1986), similar to *IRF1* mRNA and protein (Stevens & Yu-Lee, 1992; Schwartz *et al*, 2023).

Interestingly, our screen specifically identified the essential exon junction complex (EJC) factors *RBM8A*, *MAGOH*, and *EIF4A3* as strong and specific positive regulators of IFNγ-induced IRF1 after 2.5 days of dox induction (Fig. 1C,D, and Fig. EV1B). Furthermore, multiple splicing and export factors that cooperate with the EJC, including *SRSF1* and *ALYREF*, scored as hits in our screen (Fig. 1D). In contrast, factors involved in other EJC-related processes, including *CASC3* and *UPF1* which are connected to translation-coupled nonsense-mediated decay (NMD), were not identified as hits in the screen (Fig. 1D).

In addition to these well-known splicing and export factors, the enigmatic RNA regulatory factor ERH was identified as a strong determinant of IFNγ-induced IRF1 expression at day 5 post-Cas9 induction (Fig. 1C,D and Fig. EV1B). ERH is not reported to have nucleic acid binding capacities but is instead thought to assist in forming protein complexes in diverse RNA regulatory processes (Kozlowski, 2023). Notably, it has been linked to the EJC (Singh *et al*, 2012), pre-mRNA splicing (Weng *et al*, 2012; Kavanaugh *et al*, 2015; Weng *et al*, 2015), and RNA export (Pacheco-Fiallos *et al*, 2023; Shcherbakova *et al*, 2023; Dufu *et al*, 2010). Consistent with a role as a facilitating factor, it has also been reported to control other RNA-related processes in cells, including piRNA processing (Zeng *et al*, 2019; Perez-Borrajero *et al*, 2021), miRNA maturation (Kwon *et al*, 2020; Fang & Bartel, 2020), transcription (Amente *et al*, 2005; Kwak *et al*, 2003), and repressive heterochromatin formation (McCarthy *et al*, 2021).

ERH and the EJC are involved at multiple steps in post-transcriptional mRNA regulation and the EJC is loaded onto all spliced mRNAs (Schlautmann & Gehring, 2020; Obrdlik *et al*, 2019; Saulière *et al*, 2012). Therefore, the possibility of a specific role in IFNγ-dependent IRF1, but not MYC, expression was surprising (Fig. 1D). To confirm this specificity, the core EJC component *MAGOH*, the post-transcriptional integrator *ERH*, and the positive control *JAK2* were knocked out individually, and intracellular IRF1 and MYC levels were determined by FACS (Fig. 1E). Consistent with the screens, inducible *MAGOH* and *ERH* knockouts reduced IFNγ-induced IRF1 levels without affecting MYC under comparable experimental conditions (Fig. 1E). From these results we concluded that EJC- and ERH-related splicing and mRNA export are more critical for IFNγ-induced expression of ISGs, such as IRF1, than for global mRNA regulation of other factors like MYC.

Given the functional association of the EJC and ERH with RNA processing, we hypothesized that *IRF1* levels were transcriptionally or post-transcriptionally regulated. To test this, we used RT-qPCR to investigate whether ERH and MAGOH regulated *IRF1* mRNA levels, and whether this was dependent on IFNγ.

*ERH* or *MAGOH* ablation significantly reduced IFNγ-induced *IRF1* mRNA levels, but not *IRF1* baseline levels (Fig. 1F). Furthermore, ERH depletion similarly reduced *IRF1* pre-mRNA levels (Fig. EV1C) yet did not affect *IRF1* mRNA turnover as determined by actinomycin D chase and RT-qPCR analysis (Fig. EV1D). Thus, our data indicate that loss of *ERH* or *EJC* components specifically reduces IFNγ-dependent signaling and consequently prevents the transcriptional induction of ISGs such as IRF1.

### ERH maintains *JAK2* mRNA levels, and consequently, IFNγ signaling

Since ERH was critical for IFNγ-induced *IRF1* transcription, we hypothesized that ERH regulates IFNγ signaling and hence global ISG expression. Moreover, as ERH is highly conserved (Wojcik *et al*, 1994) we reasoned that its role in IFNγ-induced transcription would also be conserved across cell types and species. To investigate whether loss of *ERH* abolishes ISG induction across species and cell types, we performed 3’ mRNA QuantSeq in IFNγ-treated or non-treated RKO-iCas9 cells (human colon carcinoma) (de Almeida *et al*, 2021) and RAW 264.7-iCas9 cells (murine macrophage) (Scinicariello *et al*, 2022).

Under non-treated baseline conditions, ERH depletion affected expression of less than 5% of all genes (Source data Fig. 2A and Source data Fig. EV2A), consistent with the notion that ERH is specific and only affects a distinct group of mRNAs (Fig. 2A and Fig. EV2A). *ERH* ablation significantly affected expression (absolute fold change ≥ 2 and padj ≤ 0.05) of 809 (548 up & 261 down) genes in RKO cells, and 545 (361 up & 184 down) in RAW 264.7 cells. These changes correspond well with a previous study that reported 785 (526 up & 262 down) or 422 (243 up and 179 down) mRNA changes after siRNA-mediated *ERH* knockdown in non-differentiated or differentiated human fibroblasts (McCarthy *et al*, 2021). In both RKO and RAW 264.7 cells, *ERH* ablation reduced expression of multiple mRNAs involved in DNA replication and DNA repair (Fig. EV2B), consistent with previous observations following ERH depletion (McCarthy *et al*, 2021; Kavanaugh *et al*, 2015).

**Figure 2.**
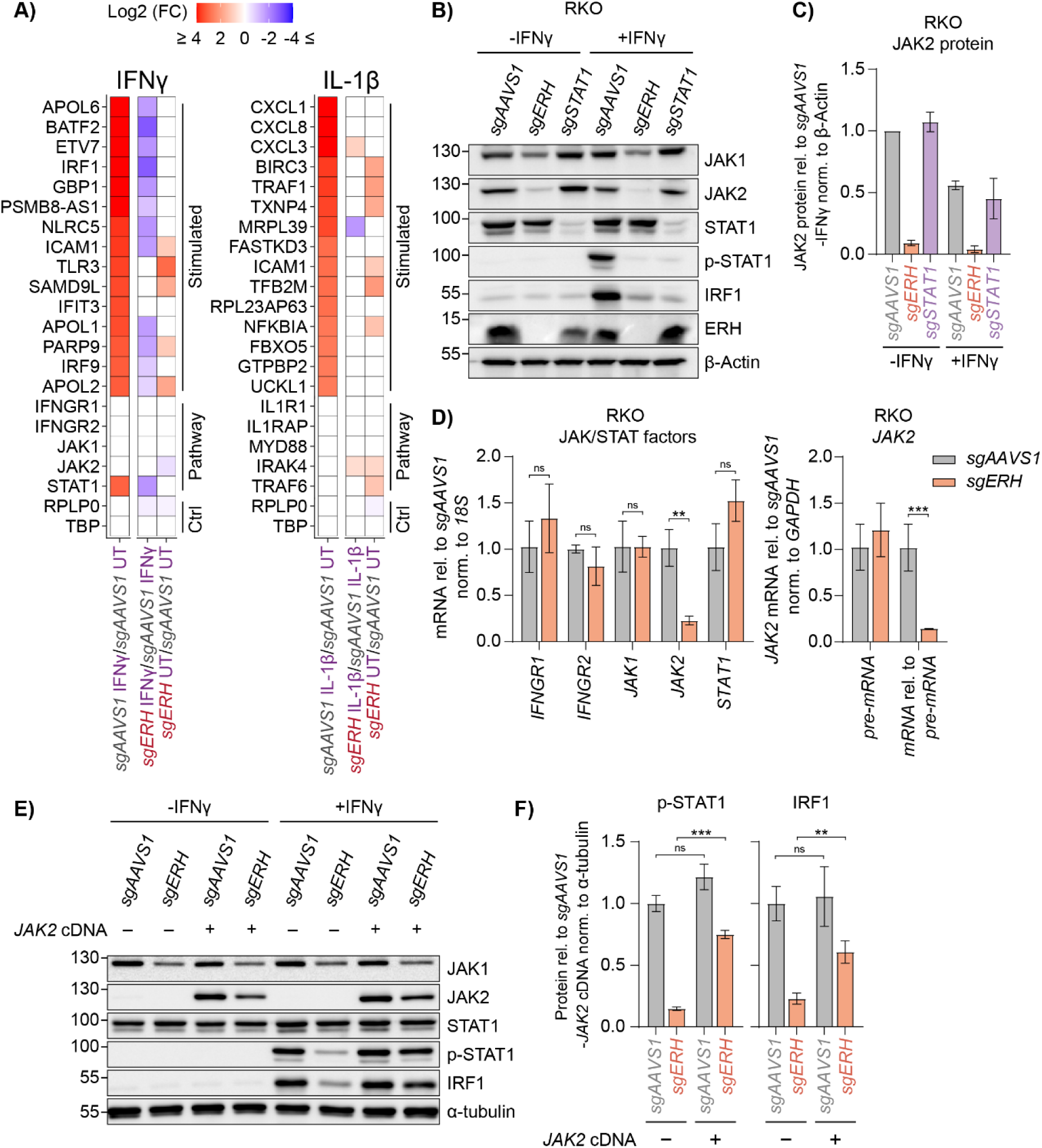
ERH is essential for maintaining *JAK2* mRNA and JAK2 protein levels, and consequently critical for IFNγ signaling. **(A)** RKO-iCas9 cells expressing the indicated sgRNAs were treated with dox for 5 days to induce Cas9 editing. The cells were then stimulated with IFNγ or IL-1β, after which RNA levels were quantified by 3’ QuantSeq. The top 15 stimulated genes and pathway components are shown. Non-significant changes (padj ≤ 0.1) were given a 0-fold change difference. n = 3 or n = 2 biological replicates; replicate outliers were excluded from the analysis. (**B**) Dox-induced RKO-iCas9 cells expressing the indicated sgRNAs were analyzed by WB, and (**C**) quantified. Data represent means and sd; n = 2 biological replicates. (**D**) RKO-iCas9 cells with the indicated genes targeted for 5 days were analyzed by RT-qPCR. Data represent means and sd; n = 3 biological replicates. Two-tailed t-test with Benjamini-Hochberg correction (*p ≤ 0.05; **p ≤ 0.01; ***p ≤ 0.001; ****p ≤ 0.0001). (**E**) RKO-iCas9 cells expressing the indicated sgRNAs were transduced with a *JAK2* cDNA expressing vector, stimulated with IFNγ for 4 h, after which protein lysates were analyzed by WB, and (**F**) quantified. Data represent means and sd; n = 3 biological replicates. Unpaired t-test with Welch correction and with Benjamini-Hochberg correction (*p ≤ 0.05; **p ≤ 0.01; ***p ≤ 0.001; ****p ≤ 0.0001).

As predicted, *ERH* ablation specifically abolished IFNγ-induced but not baseline mRNA levels of ISGs, including *IRF1* and *BATF2*, in RKO cells (Fig. 2A), indicating that ERH is indeed crucial for IFNγ-induced ISG transcription. Likewise, in RAW 264.7 cells, *ERH* ablation significantly reduced IFNγ-induced expression of ISGs, including *Irf1*, *Batf2*, and *Cxcl10* (Fig. EV2A). To assess whether ERH was specifically important for IFNγ signaling or has a more general effect on rapid cytokine-induced transcription of response genes, we treated cells in parallel with IL-1β as an alternative cytokine stimulus. In contrast to IFNγ-induced ISGs, ERH depletion did not reduce IL-1β-induced mRNA expression of response genes such as CXCL factors in neither RKO (Fig. 2A), nor RAW 264.7 cells (Fig. EV2A). In summary, ERH plays a highly specific regulatory role in IFNγ-induced gene expression across cell types and species.

Given the specific role of ERH in controlling the type II IFN axis, we reasoned that ERH may regulate the levels of JAK/STAT factors, which are essential for IFNγ signaling (Fig. 1A). To systematically test how *ERH* ablation impacts JAK/STAT pathway components, their protein levels were analyzed by western blot (WB) in lysates from sg*ERH* cells. Loss of *ERH* strongly reduced STAT1 activation, as evidenced by its reduced phosphorylation upon IFNγ treatment, without major effects on total STAT1 levels (Fig. 2B). This indicated that the effect of *ERH* loss lies upstream of STAT1. In line with this hypothesis, *ERH* targeting led to an almost complete loss of JAK2 under both stimulated and unstimulated conditions (Fig. 2B,C), arguing that ERH maintains baseline JAK2 levels.

Given that ERH functions in various RNA regulatory processes, we investigated whether reduced *JAK2* mRNA levels explained the loss of JAK2 protein. *ERH* ablation reduced *JAK2* mRNA baseline levels in 3’ mRNA QuantSeq data without affecting the expression of other JAK/STAT factors (Fig. 2A and Fig. EV2A). RT-qPCR measurements of mRNAs encoding the individual JAK/STAT pathway components (IFNGR1, IFNGR2, JAK1, JAK2, and STAT1) further confirmed that *ERH* knockout exclusively reduced *JAK2* transcript levels, which were decreased by approximately 10-fold (Fig. 2D).

Next, we examined *JAK2* pre-mRNA levels using intron-exon junction primers. While *ERH* ablation reduced mature *JAK2* mRNA levels, it did not significantly affect *JAK2* pre-mRNA levels (Fig. 2D). Thus, ERH is important for post-transcriptional *JAK2* processing yet has minimal effects on *JAK2* transcription. These effects were also conserved in RAW 264.7 cells, as *ERH* ablation similarly reduced JAK2 protein levels (Fig. EV2C,D) and *JAK2* mature-mRNA levels relative to pre-mRNA in these cells (Fig. EV2E). The effect size in RAW 264.7 was lower than in RKO cells, consistent with less efficient ERH depletion after 5 days of Cas9-induced editing (Fig. 2B and Fig. EV2C).

To test whether the reduced JAK2 levels caused abrogated IFNγ signaling in *ERH*-targeted cells, we rescued *JAK2* expression in these cells by inserting a cDNA JAK2-P2A-mCherry expression vector (Fig. EV2F). As predicted, exogenous expression of JAK2 largely rescued JAK/STAT signaling and IFNγ-driven ISG transcription in *ERH* KO cells (Fig. 2E) as determined by significantly increased p-STAT1 and IRF1 levels (5-fold increase, padj ≤ 0.001, and 2.7-fold increase, padj ≤ 0.01, respectively) (Fig. 2F).

Taken together, our data show that ERH has a conserved role in post-transcriptional processing of *JAK2* mRNA under homeostatic conditions, and consequently is critical for IFNγ-induced JAK/STAT-driven ISG expression.

### ERH associates with the EJC and EJC-associated splicing and export factors that are critical for JAK2 production

To elucidate the function of ERH, we analyzed its associated proteins. To this end, we exogenously expressed ERH-mTurquoise2 in HEK-293T cells and employed co-IP tandem mass spectrometry (MS/MS) (Fig. EV3A). As expected, ERH itself and its previously described direct protein-protein interaction partners such as POLDIP3 (Smyk *et al*, 2006) were significantly enriched in the pull-down (Fig. 3A), validating our experimental setup.

**Figure 3.**
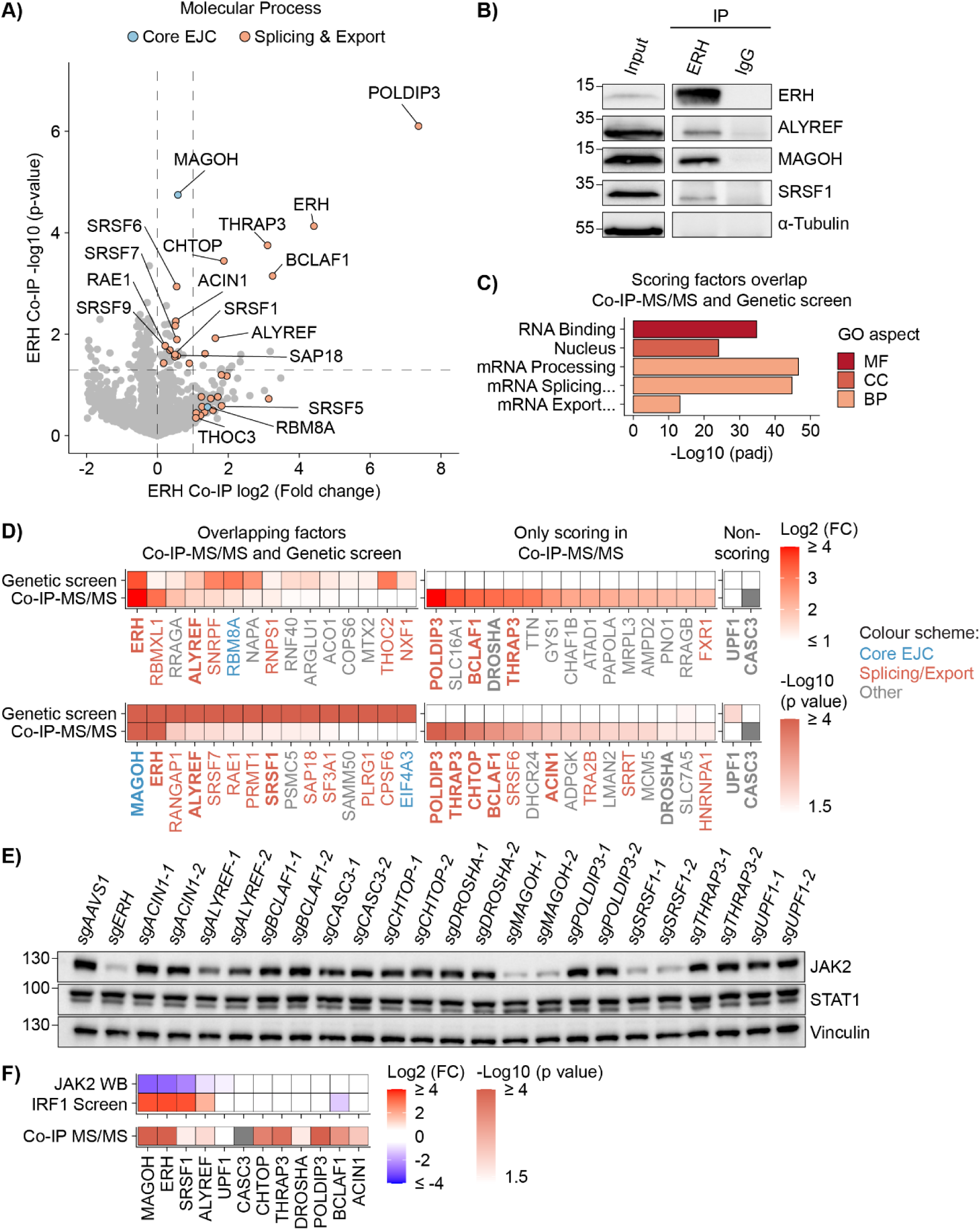
EJC-associated splicing and export factors interact with ERH and are critical for JAK2 production. **(A)** ERH-mTurqoise2 was isolated from HEK-293T cells by immunoprecipitation (IP), and interacting proteins identified by tandem mass spectrometry (MS/MS). Cells not expressing the ERH-mTurqoise2 construct were used as comparative controls. Dashed lines indicate enriched proteins based on adjusted p value (p ≤ 0.05 and log2 fold change > 0), or fold change (log2 fold change ≥ 1). n = 3 biological replicates. (**B**) Endogenous ERH IP from RKO-iCas9 cells, and indicated interactors analyzed by WB. Representative immunoblot of n = 2 biological replicates. (**C**) Selected top GO terms for overlapping genes enriched in ERH co-IP-MS/MS and the genetic screen of IFNγ signaling regulators. MF, molecular function; CC, cellular component; BP, biological process. (**D**) Heatmaps showing the top 15 enriched proteins in ERH co-IP-MS/MS based on fold change or adjusted p value, that either scored (left) or did not score (right) in the genetic screen for IFNγ regulators. For each factor, the timepoint post Cas9 induction with lowest p value or highest fold change from the genetic screen is shown. Factors are color-coded for involvement in: EJC (blue), splicing and export (orange), or other processes (gray). Non-scoring cytoplasmic factors and other factors used for validation in panel (**E**) are indicated in bold. (**E**) RKO-iCas9 cells with the indicated genes targeted for 5 days were analyzed for JAK2 and STAT1 protein levels by WB. Two different sgRNAs targeting each gene are indicated with “1” or “2”. (**F**) Heatmaps showing from top to bottom: mean JAK2 protein levels quantified from panel (**E**), enrichment in the genetic screen for IFNγ regulators at 5 days of Cas9 induction (Fig. 1C), and adj. p value in ERH co-IP-MS/MS from panel (**A**).

Consistent with previous data showing that EJC components (Singh *et al*, 2012) and related splicing and export factors such as the TREX complex (Pacheco-Fiallos *et al*, 2023; Dufu *et al*, 2010) pull down ERH, we detected a significant enrichment of peptides for MAGOH/MAGOHB, ALYREF, and SRSF1 in our ERH pulldown (Fig. 3A). Independent co-IPs of endogenous ERH from RKO cells further confirmed these interactions (Fig. 3B). These results suggest that the EJC and its associated factors may function with ERH in the same mRNP complexes.

Several of the identified interacting factors also scored well in our genetic screen for regulators of IFNγ signaling (Fig. 1C). Factors that both physically interacted with ERH by co-IP and genetically phenocopied the effect of *ERH* loss tended to be linked to nuclear post-transcriptional RNA processing, particularly splicing and export (Fig. 3C,D – left side). Additional nuclear factors that regulate splicing and export were solely identified by co-IP MS/MS, including BCLAF1, THRAP3, and ACIN1 (Fig. EV3B and Fig. 3D – right side), consistent with the predominantly nuclear localization of endogenous ERH in RKO (Fig. EV3C) and other cells (Banko *et al*, 2013). Together, the ERH interactome and the identified IFNγ-regulatory factors support a critical role of ERH in *JAK2* post-transcriptional mRNA processing.

To determine if nuclear factors interacting with ERH also regulate *JAK2*, each factor was targeted with two independent sgRNAs, after which JAK2 protein levels were measured by WB (Fig. 3E). In addition, total STAT1 levels were analyzed as a proxy for global cellular effects caused by the knockouts. WB measurements confirmed the effective and specific depletion of ERH, MAGOH, ALYREF, and SRSF1 in targeted cells (Fig. EV3D). We chose three groups of factors to investigate based on their interaction with ERH and how they scored in the genetic screen: **i)** factors interacting by co-IP-MS/MS and scoring in the genetic screen (i.e., factors that might facilitate *JAK2* processing: ERH, ALYREF, MAGOH, and SRSF1), **ii)** factors interacting by co-IP-MS/MS but not scoring in the genetic screen (i.e., potentially cooperative, but not functionally limiting factors: ACIN1, BCLAF1, THRAP3, CHTOP, DROSHA, and POLDIP3), and **iii)** non-interacting and non-scoring factors (i.e. negative controls such as cytoplasmic RNA regulatory factors: CASC3 and UPF1).

As predicted, the first group of factors (interacting by co-IP-MS/MS and scoring well in the genetic screen), significantly and specifically reduced JAK2, but not STAT1, protein levels (Fig. 3F and Fig. EV3E). *MAGOH* and *SRSF1* ablation, and to a lesser extent *ALYREF* loss, reduced JAK2 levels similarly to *ERH* ablation (∼7-fold and ∼2 fold, respectively). In contrast, depletion of non-scoring factors (groups two and three) did not affect or only mildly reduced JAK2 (∼23% reduction), and did not affect STAT1 levels (Fig. 3F and Fig. EV3E). In conclusion, our results indicate that ERH and its interacting splicing and export factors such as MAGOH, SRSF1, and ALYREF together regulate *JAK2* mRNA processing.

### Defects in *JAK2* mRNA processing prevent its subsequent nuclear export

Since ERH interacted with nuclear mRNA processing factors, we hypothesized that *ERH* controls the nuclear/cytoplasmic mRNA localization of *JAK2* and a subset of other cellular mRNAs that are likewise regulated.

To determine how ERH influenced the relative subcellular mRNA concentrations of *JAK2* and to identify similar mRNAs that are post-transcriptionally affected, we performed subcellular fractionation and 3’ mRNA QuantSeq in RKO-iCas9 cells. To identify functional similarities and differences between ERH and its interactors, we also targeted *MAGOH*, *SRSF1*, and *ALYREF*. In addition, *POLDIP3* was targeted as a negative control, since it interacts with ERH but does not affect *JAK2* levels (Fig. 3F). Robust fractionation was confirmed by WB using the cytosolic marker tubulin and the nuclear marker Lamin A/C (Fig. EV4A). The sequencing experiments were performed without spike-ins for general mRNA abundance, and thus measured relative changes in mRNA levels.

There is extensive cross-talk between transcriptional, post-transcriptional, and mRNA degradation machineries, reviewed in (Maniatis & Reed, 2002; Rambout *et al*, 2018; Rambout & Maquat, 2024). Therefore, the knockouts could affect sub-cellular mRNA distributions that are indicative of transcriptional or post-transcriptional effects in three major ways: **i)** mRNAs with cytoplasmic depletion and increased or unaffected nuclear retention, indicative of disrupted post-transcriptional mRNA processing connected to either splicing, export, translation, or mRNA degradation by NMD, **ii)** mRNAs with increased nuclear and unaffected cytoplasmic levels, indicative of increased transcriptional output and altered post-transcriptional processing, and **iii)** mRNAs with both decreased or increased nuclear and cytoplasmic abundance, indicative of altered transcription.

Consistent with ERH having a post-transcriptional role, QuantSeq analysis of the fractions revealed that *ERH* ablation specifically reduced cytoplasmic levels and/or increased nuclear levels of multiple mRNAs (Fig. 4A). Depletion of ERH significantly reduced cytoplasmic (log2FC -1.2 and padj 7.1e^-08^) and increased nuclear (log2FC 1.2 and padj 1.9e^-08^) *JAK2* mRNA abundance (Fig. 4A), indicating that *ERH* ablation disrupted *JAK2* post-transcriptional mRNA processing, resulting in *JAK2* nuclear retention.

**Figure 4.**
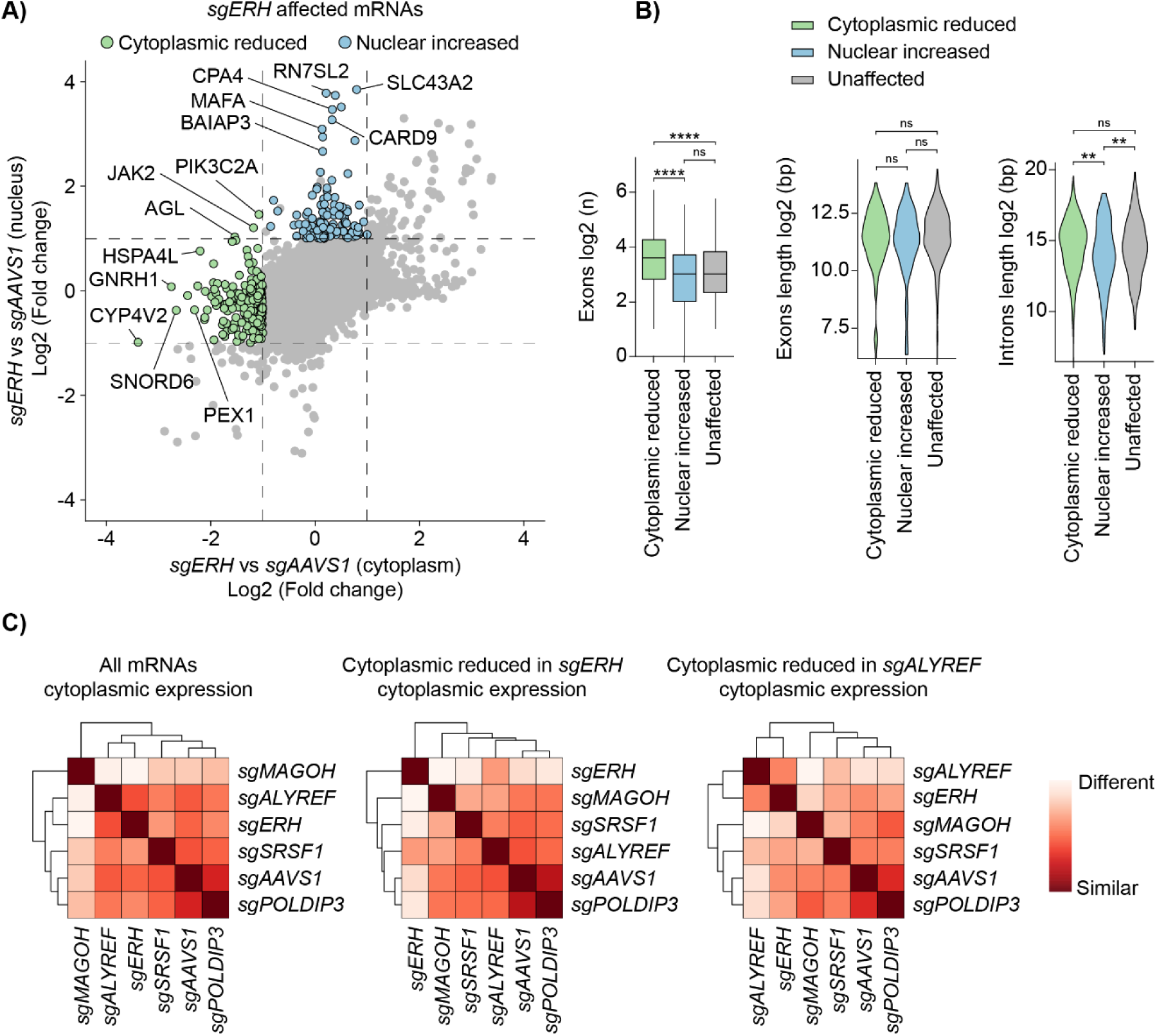
ERH prevents faulty post-transcriptional *JAK2* mRNA processing. **(A)** RKO-iCas9 cells expressing Cas9 for 5 days and sg*AAVS1* or sg*ERH* were fractionated and their mRNA levels quantified by 3’ QuantSeq. Dashed lines indicate absolute log2 fold expression change ≥ 1. n = 3 biological replicates. (**B**) Log2 quantification of number of exons (left), or length of exons (middle) and introns (right) in mRNA subsets from (A) shown by box plot or violin plots, respectively. Number of mRNAs (n) for each group from left to right: 244, 121, and randomly sampled 365. Two-sided Wilcoxon Rank Sum test (*p ≤ 0.05; **p ≤ 0.01; ***p ≤ 0.001; ****p ≤ 0.0001). (**C**) Hierarchical clustered heatmap showing the distance between RKO sample groups with indicated knockouts in subcellularly fractionated 3’ QuantSeq based on cytoplasmic variance stabilizing transformed (vst) mean mRNA expression from biological triplicates (n = 3). Subsets used for analysis from left to right: all mRNAs (n = 17657), mRNAs with reduced cytoplasmic expression in *sgERH* as in (A), n = 248, and mRNAs with reduced cytoplasmic expression in *sgALYREF*, n = 117.

Faulty transcripts retained in the nucleus can be targeted for degradation (Boutz *et al*, 2015; Davidson *et al*, 2019; Rosa-Mercado & Steitz, 2022). Therefore, we reasoned that mRNAs with considerably reduced cytosolic levels, yet minimally affected nuclear concentrations, likely also represent transcripts that are post-transcriptionally regulated by ERH. Other mRNAs that phenocopied *JAK2* and were likewise reduced in the cytosol upon *ERH* KO included multiple DNA repair and DNA replication factors (Fig. EV4B), consistent with previous reports of ERH regulating splicing of such mRNAs (Kavanaugh *et al*, 2015; Weng *et al*, 2015). In contrast to mRNAs reduced in the cytoplasm, only minor similarities and enriched GO terms were identified for mRNAs only affected in the nucleus (Fig. EV4B), suggesting that these mRNAs are regulated through a different mechanism.

To identify features that could explain the post-transcriptional effect of ERH on *JAK2*, we analyzed mRNAs that phenocopied *JAK2* behavior in *ERH* KO cells. Since *JAK2* is a distinctly large mRNA (comprised of 28 exons, and a 7 kb mature mRNA derived from a 146 kb gene), we analyzed the number of exons and length of mRNAs/genes of other candidate ERH-targets. Interestingly, mRNAs depleted in the cytoplasm, but not mRNAs accumulating in the nucleus, had significantly more exons and introns than non-affected mRNAs (Fig. 4B). However, these mRNAs and their corresponding genes were not significantly longer than average (Fig. 4B), suggesting that the number but not the length of exons/introns was an important feature for ERH-regulated mRNAs. We reasoned that a higher number of exons/introns could increase the cumulative probability for mis-splicing and result in non-exported transcripts. Taken together, these data indicated that mRNAs that decreased in the cytoplasm, but not those increased in the nucleus, exhibited *JAK2*-like features, supporting a model of altered splicing and subsequent decreased export upon ERH depletion.

Next, we compared how ERH affected overall subcellular mRNA localization compared to its interactors (MAGOH, SRSF1, ALYREF, and POLDIP3). Interestingly, distance heatmaps (Fig. 4C) and unsupervised principal component analysis (PCA) (Fig. EV4C) showed that ERH depletion affected cytoplasmic mRNA expression most similar to loss of the RNA export factor *ALYREF*. Furthermore, when comparing subsets of mRNAs with reduced cytoplasmic expression in either knockout, *ERH* and *ALYREF* ablations behaved most similarly to one another (Fig. 4C). In contrast to cytoplasmic transcript levels, nuclear affected mRNAs differed markedly between the two knockouts (Fig. EV4C,D). Consistent with a general role in mRNA export, *ALYREF* knockout resulted in strong nuclear accumulation of nearly 900 mRNAs. In comparison, less than 150 mRNAs accumulated in the nuclear fractions of *sgERH* cells. In conclusion, our results suggest that ERH directly or indirectly, for example by regulating splicing, facilitates export of a subset of mRNAs.

### ERH is critical to prevent intron retention and subsequent export disruption of *JAK2* mRNA

Our data so far indicated that ERH is critical for proper accumulation of cytosolic *JAK2* mRNA. We thus hypothesized that ERH plays a functional role in either: **i)** stabilizing cytosolic *JAK2* mRNA by preventing NMD, **ii)** exporting *JAK2* mRNA to the cytoplasm, or **iii)** proper splicing of *JAK2* mRNA, as a prerequisite for its subsequent nuclear export.

To determine which of these processes was affected in the absence of *ERH*, RNA-Seq was performed on fractionated and poly(A)-enriched RNA from sg*ERH* RKO-iCas9 cells. Parallel samples treated with the translation inhibitor cycloheximide (CHX) were included to determine if translation-dependent NMD plays a role in *JAK2* mRNA regulation by ERH. Changes in relative cytoplasmic and nuclear mRNA levels correlated well between the poly(A)-enriched RNA-Seq samples and our previous fractionated 3’ mRNA QuantSeq data, underpinning the reproducibility and comparability of the two experiments (Fig. EV5A).

To assess whether *ERH* loss resulted in mis-splicing and IR, we utilized IRFinder and DEseq2 (Middleton *et al*, 2017; Lorenzi *et al*, 2021) for differential enrichment analysis of IR in *sgERH* samples compared to the *sgAAVS1*-targeted controls. Consistent with being required for proper *JAK2* splicing, *ERH* ablation caused significant IR in nuclear *JAK2* mRNA (introns 9-14 and intron 16) and in other mRNAs with reduced cytoplasmic expression in the absence of *ERH* (Fig. 5A,B and Fig. EV5B), hence referred to as ‘*JAK2*-like’ mRNAs. Approximately 28% of these mRNAs had at least one increased IR event (in the nucleus or cytoplasm) in comparison to 3% in non-affected mRNAs (Fig. 5C). This was likewise the case for mRNAs with concurrently reduced nuclear expression (Fig. 5A), suggesting that these affected mRNAs were efficiently recognized and degraded by the nuclear RNA regulatory machinery. Moreover, multiple mRNAs with IR following *ERH* ablation were involved in DNA repair or replication (Fig. EV5C), confirming that ERH plays a role in splicing of these transcripts. In summary, ERH is required for proper splicing and prevention of IR in a specific subset of mRNAs, including *JAK2*.

**Figure 5.**
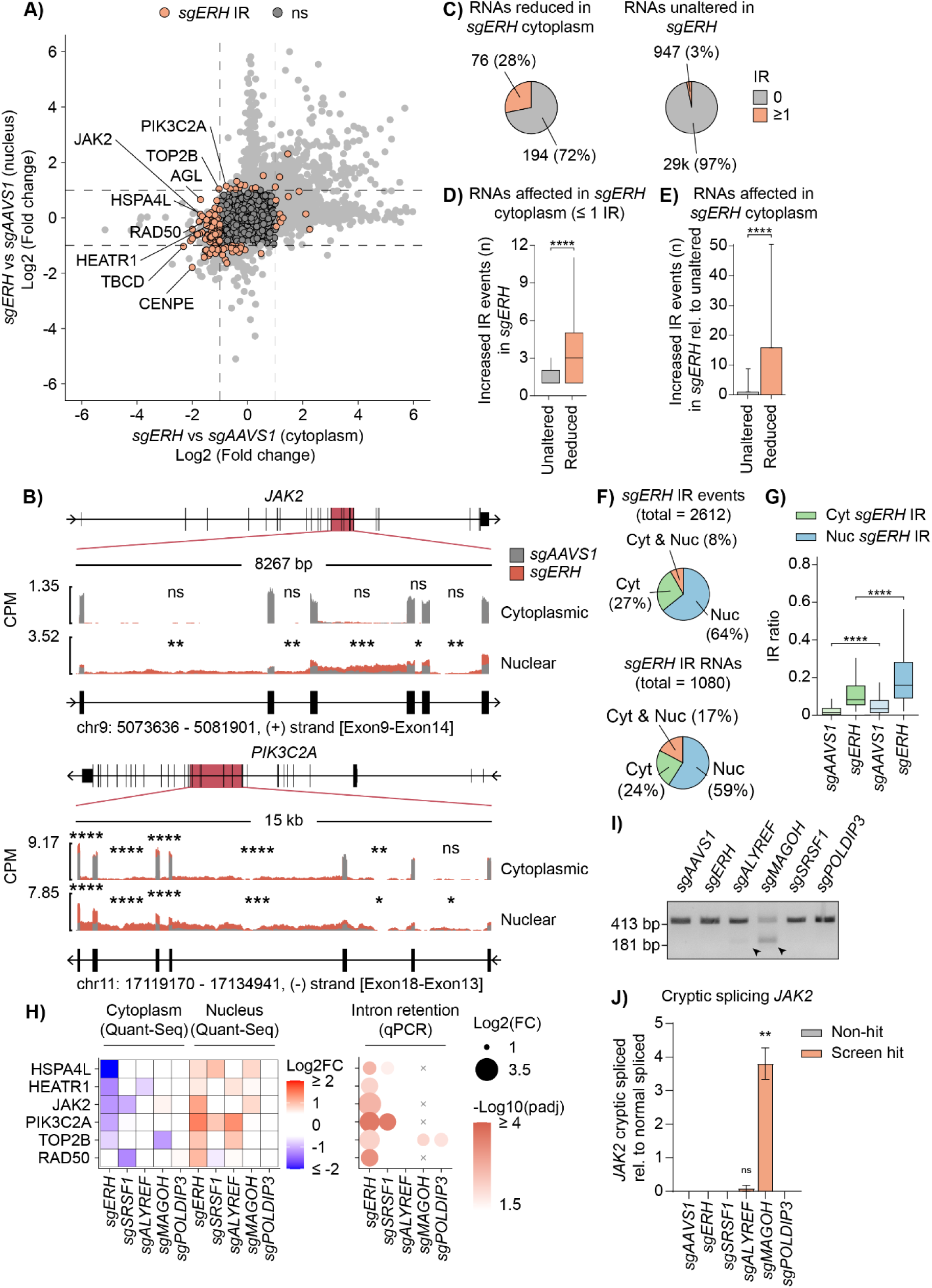
ERH prevents intron retention and disrupted export of *JAK2* and other select mRNAs. RKO-iCas9 cells with the indicated knockouts after 5 days of Cas9 induction were fractionated and subjected to poly(A)-enriched RNA-Seq analysis. (**A**) mRNAs with expression difference and at least one increased intron retention (IR) event (padj ≤ 0.05 and log2 fold change ≥ 1, nucleus or cytoplasm) in *sgERH* over *sgAAVS1* determined by IRFinder are highlighted. n = 3 biological replicates. (**B**) *JAK2* and *PIK3C2A* mRNA regions with IR in *sgERH*; mean reads per million (CPM) in *sgAAVS1* or *sgERH* are plotted. Total (top) and zoomed in (bottom) gene structures are displayed with horizontal lines indicating introns and blocks indicating exons. Significance based on adjusted p value for increased IR in *sgERH* is indicated (*p ≤ 0.05; **p ≤ 0.01; ***p ≤ 0.001; ****p ≤ 0.0001). (**C**) Pie charts depicting the number of RNAs with at least one increased IR event regulated by ERH, comparing cytosolically reduced and unaltered RNAs. (**D**) Box plots comparing number of increased IR events in *sgERH* between RNA subsets containing at least 1 increased *sgERH-*regulated IR event. Two-sided Wilcoxon Rank Sum test (*p ≤ 0.05; **p ≤ 0.01; ***p ≤ 0.001; ****p ≤ 0.0001). (**E**) Bar graph comparing the number of increased IR events in *sgERH* between RNA subsets relative to unaltered RNAs. Data represent means and sd. Two-sided Wilcoxon Rank Sum test (*p ≤ 0.05; **p ≤ 0.01; ***p ≤ 0.001; ****p ≤ 0.0001). (**F**) Pie charts depicting number of retained introns (top) or their corresponding RNAs (bottom) caused by *ERH* ablation (mean IR ratio ≥ 0.05, padj ≤ 0.05 and log2 fold change ≥ 1 between knockouts) in the cytoplasm, nucleus, or both. (**G**) Boxplots comparing mean IR ratio of retained introns in RKO-iCas9 cells with indicated knockouts. Two-sided Wilcoxon Rank Sum test with Holm-Bonferroni correction (*p ≤ 0.05; **p ≤ 0.01; ***p ≤ 0.001; ****p ≤ 0.0001). (**H**) Heatmaps of subcellularly fractionated 3’ QuantSeq (left), and intron retention measured by RT-qPCR in the same RNA samples (right.) (n = 3 biological replicates. x; not measured). (**I**) RKO-iCas9 cells expressing the indicated sgRNAs were treated with dox for 5 days to induce Cas9 expression. Total RNA was isolated and subjected to *JAK2* splicing analysis by end-point RT-PCR and agarose gel electrophoresis, and **(F)** bands were quantified. Cryptically spliced *JAK2* mRNA (181 bp amplicon) and correctly spliced *JAK2* mRNA (413 bp amplicon) are indicated. Data represent means and sd; n = 3 biological replicates. Welch’s two-tailed t-test (*p ≤ 0.05; **p ≤ 0.01; ***p ≤ 0.001; ****p ≤ 0.0001).

*ERH* ablation often caused multiple IR events in ‘*JAK2-*like’ transcripts. These transcripts had a median of three IR events compared to one in unaffected mRNAs (Fig. 5D). Taking these factors into account, cytosolically depleted mRNAs had on average fifteen times more ERH-dependent IR events compared to unaffected mRNAs (Fig. 5E). Moreover, many ‘*JAK2*-like’ transcripts, such as *PIK3C2A*, retained multiple adjacent introns in *sgERH* samples (Fig. 5B and Fig. EV5B). We determined that these IR events were more often adjacent than can be predicted by random chance (Fig. EV5D), similar to what has been previously described for general IR (Middleton *et al*, 2017). It has been suggested that connected IR events could depend on their splicing factors and other RBPs (Middleton *et al*, 2017), in line with ERH being involved in multiple RNA-related processes such as splicing and determining Pol-II elongation rates (Braunschweig *et al*, 2014) resulting from opening of chromatin (McCarthy *et al*, 2021).

To determine whether NMD contributed to reduced cytoplasmic mRNA levels in intron-retained mRNAs following *ERH* ablation, we compared non-treated and CHX treated samples. CHX treatment was confirmed to effectively inhibit NMD by the increase of *GADD45A* (Fig. EV5E), an mRNA that is constantly turned-over by NMD (Nelson *et al*; Zhao *et al*, 2020). In contrast, CHX treatment stabilized only a small fraction of the ERH-regulated cytosolically depleted mRNAs with IR (Fig. EV5E) and did not significantly increase cytoplasmic IR in these mRNAs. Thus, ERH-regulated mRNAs are not reduced by cytoplasmic control mechanisms such as NMD but undergo nuclear retention as commonly described for inefficiently spliced transcripts (Yap *et al*, 2012; de Almeida *et al*, 2010; Boutz *et al*, 2015). In line with this notion, most IR events (∼64%) caused by *ERH* ablation and their corresponding RNAs (∼59%) were only detected in the nucleus (Fig. 5F). Moreover, the degree of IR, as measured by the IR ratio, caused by *ERH* ablation was significantly higher in nuclear IR events compared to cytoplasmic IR events (median IR ratio 0.01 or 0.08 in cytoplasm, and 0.03 or.16 in nucleus, in *sgAAVS1* or *sgERH*, respectively) (Fig. 5G). In conclusion, our results indicate that in the absence of *ERH*, most ‘*JAK2*-like’ mRNAs are not exported and consequently retained in the nucleus.

To determine if ERH-interacting factors identified by co-IP MS/MS (Fig. 3A) regulate IR similarly to ERH, we analyzed by RT-qPCR whether their knockouts affected IR in the same nuclear RNA samples analyzed by 3’ mRNA QuantSeq (Fig. 4C). We analyzed six different mRNAs with the strongest ERH-regulated IR events (including *JAK2*) that had altered cytoplasmic or nuclear mRNA levels in at least one other knockout (as measured by 3’ mRNA QuantSeq) (Fig. 5H).

Consistent with our poly(A)-enriched RNA-Seq results, *ERH* ablation caused IR in all selected mRNAs, including *JAK2*. In contrast, none of the other knockouts caused IR in *JAK2*, suggesting that these factors likely act in different RNA processing steps of *JAK2* and similarly affected ‘*JAK2*-like’ mRNAs. Notably, only *SRSF1* ablation reduced cytoplasmic *JAK2* like in the *ERH* KO, however, we could not measure similar IR in the *SRSF1* KO. Interestingly, *SRSF1* ablation did show IR in mRNAs with nuclear retention, namely *PIK3C2A* and to a lesser extent *HSPA4L* (Fig. 5H), suggesting that a smaller subset of the ERH-regulated splicing events could be accomplished together with SRSF1.

We further compared the knockouts by analyzing the mean sample distance based on mRNA expression levels for mRNAs with or without ERH-regulated IR. In line with these factors likely acting at different RNA processing steps, the similarity between the knockouts did not increase when comparing mRNAs with ERH-regulated IR to those without (Fig. EV5F).

Previously, we confirmed that loss of ERH or ERH-interacting factors like *MAGOH* and *ALYREF* reduces JAK2 protein levels (Fig. 3F). Therefore, we hypothesized that these factors regulate different steps of *JAK2* post-transcriptional processing, yet -if this is a rate-limiting step-ultimately affect JAK2 protein levels in a similar way.

Knockdown of the core EJC factor *EIF4A3* has previously been shown to cause mis-splicing of cryptic splice sites as analyzed by total RNA-Seq (Schlautmann *et al*, 2022). This faulty splicing could result in non-functional and degraded proteins and could potentially explain lower JAK2 protein levels in *MAGOH* knockout cells. Indeed, erroneous splicing in exon3 of *JAK2* in the absence of EJC components has been previously reported (Schlautmann *et al*, 2022). We further validated that *MAGOH* ablation caused faulty splicing of *JAK2* by RT-qPCR (Fig. 5I), where nearly four times more mis-spliced *JAK2* was present than normally spliced *JAK2* (Fig. 5J). Similar splicing changes were also detected in the *ALYREF* knockout, which had lower (∼10%) faulty spliced *JAK2* levels (Fig. EV4F). This reduced effect is in line with *ALYREF* loss having a smaller effect on JAK2 protein and IFNγ-induced IRF1 levels (Fig. 3F).

From these results we conclude that ERH-interacting factors do not affect IR in ‘*JAK2*-like’ transcripts as ERH does, but are rather critical for different post-transcriptional steps of JAK2 mRNA processing, and in this way affect JAK2 protein and IFNγ signaling.

### ERH-regulated retained introns are AU-rich

To determine if ERH-regulated IR events had unique characteristics, we analyzed four characteristics known to influence IR (Petrova *et al*, 2022; Braunschweig *et al*, 2014): **i)** lengths of the introns and flanking exons, **ii)** 3’ and 5’ splice site strengths, **iii)** intronic and exonic GC content, and **iv)** motif-abundances for RBPs. We compared retained introns caused by *ERH* ablation with a similar number of randomly sampled introns that were similarly retained in both *sgAAVS1* and *sgERH* under homeostatic conditions, hence referred to as general IR, and with non-retained introns.

We determined that ERH-regulated introns were shorter (Fig. 6A) and had weaker 3’ splice site strength (Fig. 6B) than non-retained introns, similar to general IR and as previously reported for regularly retained introns (Petrova *et al*, 2022; Braunschweig *et al*, 2014). Interestingly, the 5’ splice site was not weaker in ERH-regulated introns whereas the 3’ splice site was even weaker than generally retained introns (Fig. 6B), suggesting that the 3’ splice site predisposes introns to ERH-mediated regulation. Furthermore, ERH-regulated introns had longer flanking exons than other introns, while total gene length was comparable to non-retained ones (Fig. EV6A). From these data we concluded that ERH-regulated introns have multiple features that have been previously reported to predispose introns for retention, such as weaker 3’ splice sites and shorter intron length.

**Figure 6.**
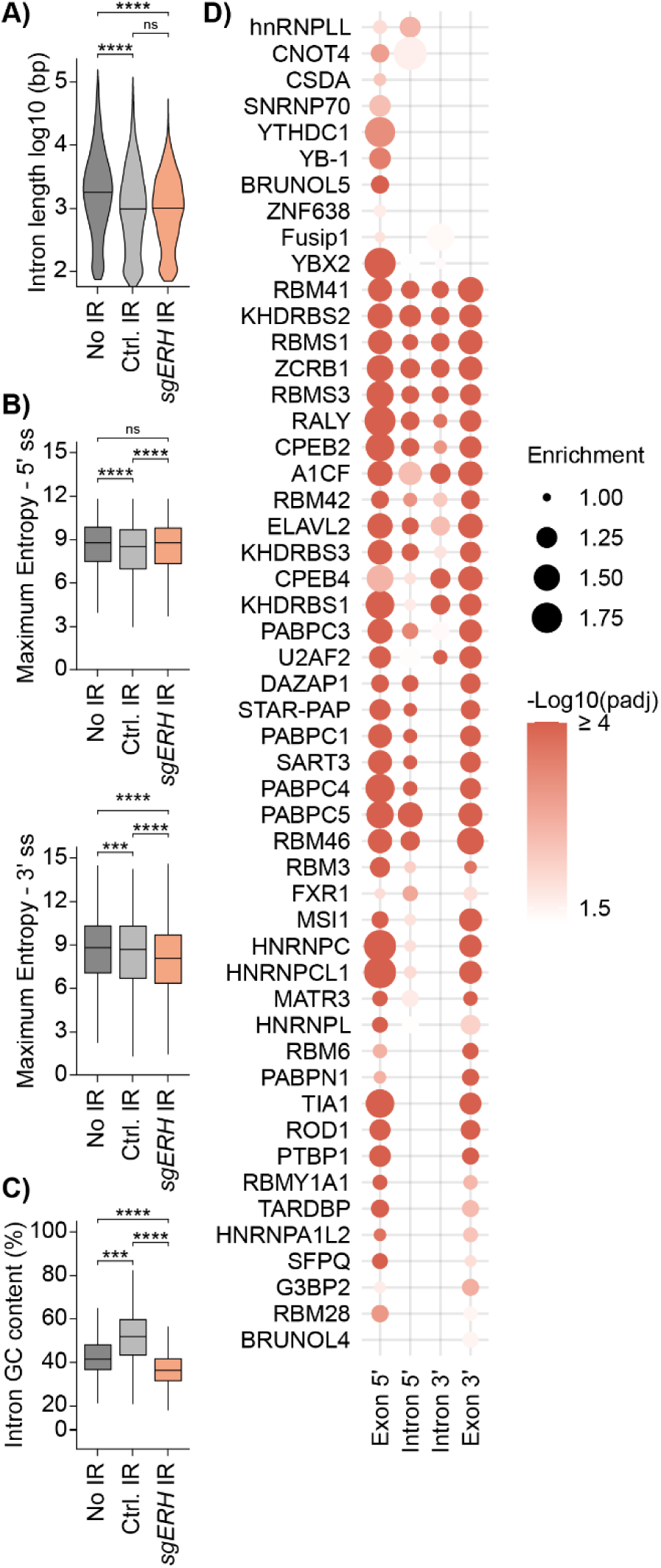
ERH-regulated retained introns and flanking exons are AU-rich. RKO-iCas9 cells with the indicated knockouts were fractionated and subjected to poly(A)-enriched RNA-Seq analysis. (**A**) Violin plots comparing length distributions between non-retained introns (“No IR”; mean IR ratio ≤ 0.05 in both *sgAAVS1* and *sgERH*), retained introns in both knockouts (“Ctrl. IR”; mean IR ratio ≥ 0.1 and absolute log2 fold change ≤ 0.2 between knockouts), or retained introns caused by *ERH* ablation (“*sgERH* IR”, mean IR ratio ≥ 0.05, padj ≤0.05 and log2 fold change ≥1 between knockouts). n = 2178 in each group (randomly sampled for control groups). (**B**) Box plots comparing average 5′ and 3′ splice site strength measured by Maximum Entropy Score (Eng *et al*, 2004). (**C**) Box plots comparing GC content. (**A-C**) Two-sided Wilcoxon Rank Sum test with Holm-Bonferroni correction (*p ≤0.05; **p ≤ 0.01; ***p ≤ 0.001; ****p ≤ 0.0001). (**D**) Heatmap of enriched (padj ≤ 0.05) RNA binding protein (RBP) motifs located near ERH-regulated retained introns compared to non-retained introns measured by simple enrichment analysis (SEA). 5′ and 3′ boundaries of flanking exons (100 bp) and introns (200 bp), excluding splice sites, were analyzed. For each RBP, the motif with the lowest padj is shown.

Previously, several types of retained introns that exhibit different features have been proposed (Braunschweig *et al*, 2014). However, all retained intron types have been reported to contain increased GC content in comparison to their non-retained counterparts (Petrova *et al*, 2022; Braunschweig *et al*, 2014). Strikingly, ERH-regulated introns (Fig. 6C) and their flanking exons (Fig. EV6B) had significantly reduced, rather than increased, GC content compared to non-retained introns, which was to a lesser extent also identified for their corresponding genes (Fig. EV6B). In line with lower GC content, ERH-dependent introns, and to an even greater degree their flanking exons, were enriched for AU-rich motifs compared to non-retained ones (Fig. 6D). The AU-rich element-binding proteins (AUBPs) linked to these motifs included multiple hnRNPs, KHDRBSs, and other RNA processing factors primarily involved in splicing, but also RNA export, supporting a regulatory role of ERH in splicing of AU-rich transcripts. This is markedly different from previous reports on regularly retained GC-rich introns that have an enrichment of SR-protein binding sites (Middleton *et al*, 2017).

In summary, our study identified ERH as a functionally critical factor for correct splicing of a select set of mRNAs, including *JAK2*. Loss of ERH-associated factors implemented in post-transcriptional processing, such as SRSF1, MAGOH, and ALYREF, likewise disrupted *JAK2* mRNA maturation. Our data indicate that these factors affect *JAK2* at different post-transcriptional steps, yet ultimately lead to the same outcome of reduced JAK2 protein levels. These findings indicate that ERH only functionally overlaps with the global splicing and export machineries for a smaller set of transcripts, which critically rely on ERH to prevent IR. Specifically, our results indicate that ERH is required for efficient splicing of introns in AU-rich regions, which differs from previously described regular GC-rich IR. Together, these findings position ERH and post-transcriptional processing as critical regulatory factors for mounting effective innate immune responses by type II IFN signaling.

## Discussion

In recent years, there have been multiple examples for genome-wide CRISPR screens that use inducible Cas9 expression to successfully capture regulatory factors regardless of their impact on cellular viability (Schwartz *et al*, 2023; de Almeida *et al*, 2021; Scinicariello *et al*, 2022). We applied this approach to identify novel positive regulatory factors of IFNγ signaling, aiming to complement previous gene-trap screens that could not identify essential factors (Brockmann *et al*, 2017). In this study, we successfully identified multiple cell-essential post-transcriptional RNA regulatory factors, namely ERH and other EJC-related splicing and export factors, as novel and specific regulators of IFNγ-induced ISG expression. Overall, we determined that these factors were critical for expression of the signaling kinase JAK2, and in this way regulated overall IFNγ-induced JAK/STAT signaling and subsequent ISG transcription.

To determine the mechanism of *JAK2* regulation we found that ERH was critical to prevent IR in *JAK2* mRNA and a subset of other similarly ERH-dependent transcripts. A significant fraction of these cytosolically reduced mRNAs could be explained by an increase in IR in *ERH* KO cells. Interestingly, the interactors of ERH investigated in this study were important for different steps of *JAK2* processing. MAGOH, and to a lesser extent ALYREF, regulated separate cryptic splicing events. Similarly, SRSF1 regulated seemingly other steps in *JAK2* mRNA processing, potentially through different IR events (Middleton *et al*, 2017) or by acting as an export adaptor (Müller-McNicoll *et al*, 2016). Overall, this highlights post-transcriptional *JAK2* processing as a critically susceptible process.

This raises the question as to what makes *JAK2* and similar mRNAs disproportionally sensitive to disruption of core RNA processing, such as splicing and export. It is clear from this study and previous ones that ablation of general acting factors, like core EJC factors or the TREX export adaptor ALYREF, disproportionally affects mRNA abundance of a subset of transcripts (Xue *et al*, 2023a; Ihashi *et al*, 2023; Schlautmann *et al*, 2022). One potential explanation for this could be the lack of compensatory and redundant mechanisms for processing of these mRNAs. Moreover, splicing and export could be relatively more important and potentially rate-limiting compared to other RNA processes such as transcription for these mRNAs. Overall, the complex interplay between different RNA processes and how this affects specific mRNAs presents an exciting avenue for future research. In particular, exploration of the molecular mechanisms by which individual RNA processing factors affect subsets of transcripts will be of high interest.

ERH acts by bridging proteins and aids in protein complex formation (Kozlowski, 2023). Thus, understanding the functional interactions with its binding partners, and the functional importance of these interactions for generating specific biological output, will be key for understanding the molecular mechanisms by which ERH regulates various cellular processes. We and other groups have shown that ERH interacts with multiple EJC-associated factors. Indeed, cross-linking mass spectrometry revealed close-range interactions between ERH and the export adaptors ALYREF and CHTOP (Pacheco-Fiallos *et al*, 2023) and the splicing/export factors BCLAF1 and THRAP3 (Shcherbakova *et al*, 2023).

We found that loss of *ERH* or *ALYREF* mimicked each other most closely in terms of cytoplasmic mRNA abundance, and *SRSF1* loss resulted in similar IR as ERH for some mRNAs. However, the overall effect of *ERH* loss, or loss of its interactors was not greater for intron-retained mRNAs, suggesting that ERH might affect these mRNAs through multiple means with its interactors. Indeed, our results indicate that intron-retained mRNAs were not extensively degraded in the cytoplasm, but rather poorly exported. The functional similarity between ERH and the export adaptor ALYREF may indicate that ERH has a role in both splicing and export of these mRNAs as a scaffolding/compaction factor in their RNP complexes. Further studies are required to explore the mechanism of ERH-dependent effects on mRNA export, for example by investigating non-spliced histone RNAs that depend on ALYREF (Fan *et al*, 2019), and in this way separate splicing from export-dependent outcomes. Despite a substantial decrease in mature target transcripts in the cytoplasm in the absence of *ERH*, their nuclear counterparts did not substantially increase in most experiments, indicating that the retained nuclear target transcripts were likely degraded in the nucleus.

We found that in the absence of *ERH*, multiple adjacent introns tended to be retained. This behavior has been previously reported (Middleton *et al*, 2017), yet its underlying reason has remained unclear. Moreover, introns retained in the absence of *ERH* were substantially more AU-rich than classically retained introns, a feature reported for mRNAs spliced at the nuclear periphery (Tammer *et al*, 2022). In line with this, we speculate that ERH could play a more dominant functional role in the nuclear periphery, and its absence could disproportionally affect transcripts spliced in that part of cellular three-dimensional space.

It should be noted that *ERH* ablation could potentially cause IR through secondary effects. An important factor shown to facilitate IR is increased chromatin accessibility, as this accelerates Pol II elongation rates resulting in alternative-splicing (Petrova *et al*, 2022). Previous work showed that ERH can be important to maintain repressive chromatin modifications (McCarthy *et al*, 2021), and we cannot exclude that increased chromatin accessibility by *ERH* ablation could have influenced IR. Our identified ERH-regulated IR events differed from conventional retained introns in having increased proximal AU-rich motifs with many corresponding AUBPs being involved in splicing and export, indicating that chromatin accessibility is unlikely to be a critical aspect for ERH-specificity. The fact that these factors were not identified in our co-IP-MS/MS could suggest that they are only transiently associated to the RNP complex during splicing. Some of the factors, such as HNRNPL, also scored in our genetic screen, but most did not, indicating they might act redundantly or that only some of them are functionally involved in these splicing events. Taken together, our results are in favor of a model in which ERH prevents AU-rich IR in a splicing dependent manner, thereby ensuring faithful export of mRNAs, including *JAK2*.

*ERH* is highly evolutionary conserved in eukaryotes, ranging from yeast to humans (Kozlowski, 2023). In contrast, the *JAK2* gene arose much later in evolution in jawed-vertebrates (Gu *et al*, 2002). Moreover, ERH is constitutively expressed at similar high levels across all human tissue types (Smyk *et al*, 2006), and is not IFNγ-induced (Fig. 2B). This makes it unlikely that ERH has adapted to directly control innate immunity. Instead, it suggests that intrinsic *JAK2* mRNA features make it critically dependent on ERH-dependent processing. Nevertheless, an increasing number of studies underpins the importance of post-transcriptional processing for generating immune output (Ullrich & Guigó, 2020; O’Connor *et al*, 2015; Green *et al*, 2020; Wagner *et al*, 2021). Therefore, it could be that some immune transcripts have evolved to be processed by core post-transcriptional machineries to ensure proper immune output.

In this context, some retained intronic transcripts (referred to as detained intronic transcripts) can serve as reservoirs that can be activated and spliced in response to dynamic stimuli such as cellular stress, DNA damage, and viral infection (Boutz *et al*, 2015; Mauger *et al*, 2016; Ninomiya *et al*, 2011). Such a dynamic role makes sense for ERH as the IR-regulated mRNAs are involved in innate immune signaling (*JAK2*) and the DNA damage response. However, mRNAs with conventional GC-rich detained introns are seemingly resistant to exosomal decay and accumulate in nuclear speckles (Boutz *et al*, 2015; Hett & West, 2014; Girard *et al*, 2012) – nuclear domains that play a role in retention and quality control of detained mRNAs (Wegener & Müller-McNicoll, 2018). In contrast, ERH-regulated mRNAs with AU-rich IR did not generally accumulate in the nucleus, indicating that they may be cleared by exosomal decay, as has been reported for inefficiently spliced transcripts (Yap *et al*, 2012; de Almeida *et al*, 2010). Determining whether ERH always aids in proper splicing and export of these mRNAs or if it dynamically influences IR in response to different stimuli will be an exciting avenue for future research.

## Methods

### Vectors

All plasmids and sgRNAs used in this study are listed in Table 1 and Table 2, respectively. The lentiviral human genome-wide sgRNA library (six sgRNAs/gene) and the lentiviral vectors driving the expression of single sgRNAs (U6 promoter) and eBFP2 or iRFP (PGK promoter) were described previously (de Almeida *et al*, 2021). The lentiviral JAK2 expression vector was obtained by cloning JAK2 cDNA (Addgene plasmid 23915) into a modified pLX303 vector that co-expressed myc-tagged mCherry through a P2A site. The ERH expression vector with C-terminally tagged mTurquoise2 used in co-IP-MS/MS was cloned in the pEGFP-C1 backbone (Clontech).

### Cell culture and reagents

All cell lines used in this study and their applications are listed in Table 3, reagents in Table 4, and antibodies in Table 5. Cells tested negative for mycoplasma contamination and the used cell lines in this study are not on the current International Cell Line Authentication Committee (ICLAC) list of commonly misidentified cell lines (version 13). RKO and RAW 264.7 parental cells were authenticated by short tandem repeat analysis. Dox-inducible Cas9 single-cell-derivative clones were generated previously (de Almeida *et al*, 2021; Scinicariello *et al*, 2022), and active Cas9 function was validated to solely occur in the presence of dox through knockout of essential genes and competitive proliferation assays. To obtain JAK2-expressing cells, pLX303-SFFV-myc-mCherry-P2A-JAK2 was transduced into RKO-Dox-Cas9 cells, and cells expressing mCherry were bulk sorted with a FACSAria III (BD Biosciences). All cells were cultured at 37°C and 5% CO_2_ in a humidified incubator.

### FACS-based CRISPR–Cas9 screen

The genome-wide FACS-based screen and generation of next-generation sequencing libraries were performed as previously described for RKO cells (de Almeida *et al*, 2021). In short, Cas9-mediated knockouts were induced by treatment with dox for 2.5 or 5 days, IRF1 expression was stimulated for 24 h pre-harvesting with human IFNγ at 10 ng/mL (Enzo Life Sciences, ENZ-PRT141-0100). IRF1 levels were determined through intracellular staining with a PE-conjugated anti-IRF1 antibody (Cell Signaling Technology, 12732).

CRISPR screen analysis was performed with MAGeCK (Li *et al*, 2014), as previously described (de Almeida *et al*, 2021). The workflows for quantification of raw sequencing reads (https://github.com/ZuberLab/crispr-process-nf) and enrichment/depletion analysis of sgRNAs using MAGeCK (https://github.com/ZuberLab/crispr-mageck-nf) are available online. Molecular process labelling was guided by literature research and GO term annotations accessed with AmiGO 2 version 2.5.17 (Carbon *et al*, 2009; Carbon & Mungall, 2024), for JAK/STAT pathway (GO:0060333), RNA splicing (GO:0008380), RNA export (GO:0006405), NMD (GO:0000184), EJC (GO:0035145), and type I IFN (GO:0060337).

### Lentivirus-like particle production and transduction

Semi-confluent Lenti-X cells were transfected with mixes composed of a lentiviral transfer plasmid of interest, pCRV1-Gag-Pol (Hatziioannou *et al*, 2004), and pHCMV-VSV-G (Yee *et al*, 1994), by using polyethylenimine (PEI, Polysciences, 23966) in a 1:3 ratio (μg DNA/μg PEI) in DMEM without supplements. Virus-like particle (VLP)-containing supernatant was cleared of cellular debris by filtration (0.45 μm) and were kept at 4°C or -80°C for short-term or long-term storage, respectively. Target cells were transduced in the presence of 5 μg/ml (RKO) or 6 μg/ml (RAW 264.7) of polybrene (Sigma-Aldrich, TR1003G).

### Intracellular staining for flow cytometry

Cell fixation, permeabilization, and staining were performed as previously described (Scinicariello *et al*, 2022). Permeabilized cells were blocked with Human BD Fc Block (BD Biosciences, 564220). Samples were analyzed on an LSRFortessa (BD Biosciences) with BD FACSDiva software (v8.0). FACS data were further analyzed using FlowJo (v10.8).

### Western blotting

Cells were lysed in RIPA lysis buffer (50 mM Tris-HCl pH 7.4, 150 mM NaCl, 1% SDS, 0.5% Sodium deoxycholate, 1% Triton X-100), rotated for 30 min at 4°C and centrifuged at 16,000 x g for 20 min at 4°C. Protein concentrations of the supernatants were determined using the Pierce BCA Protein Assay Kit (Thermo Fisher Scientific, 23225). Lysates were boiled at 95°C for 5 min in Laemmli sample buffer with 10% β-mercaptoethanol. Lysates (20 μg) were loaded on SDS polyacrylamide gels with varying percentages based on MW of proteins of interest. Proteins were blotted on nitrocellulose membranes at 4°C for 75 min at 300 mA in Towbin buffer (25 mM Tris-HCl pH 8.3, 192 mM glycine, and 20% ethanol). Membranes were blocked in 5% BSA in PBS-T for 1 h at room temperature, and subsequently incubated with primary antibody overnight at 4°C while shaking. Membranes were washed five times with PBS-T and incubated with HRP-coupled secondary antibody for 1 h at room temperature and imaged with the ChemiDoc Imaging System from Bio-Rad. Relative protein levels were quantified with Image Lab (BioRad) on non-saturated exposures.

### Total RNA isolation, cDNA synthesis, and qPCR

Total RNA was harvested from 1-2 x 10^6^ RKO-iCas9 or RAW 264.7-iCas9 cells by Trizol lysis (Thermo-Fisher Scientific, 5596–018) and total RNA was isolated per the manufacturer’s recommendations. The RNA was treated with Turbo DNase (Thermo Fisher Scientific, AM2238), and cDNA was prepared with Oligo dT18 primers (Thermo Fisher Scientific, S0132) or random hexamer primers (Thermo Fisher Scientific, S0142) using RevertAid Reverse Transcriptase (Thermo-Fisher Scientific, EP0441) as per the manufacturers’ recommendations. qPCR samples were analyzed in technical triplicates or duplicates and with H_2_O and no reverse transcriptase controls on a Mastercycler (Biorad) using 1x qPCR MM (10 mM Tris pH 8.5, 50 mM KCl, 0.15% Triton X-100, 2 mM MgCl2, 200 μM dNTPs (Promega, U1515), 200 mM Trehalose, 2.5% Formamide (Applichem, A2156), 0.5x SYBR Green I (Thermo-Fisher Scientific, S7567), and 25 U/mL Taq polymerase (Promega, M7484B), in nuclease free H_2_O (Ambion, AM9930)). Relative changes in gene expression were calculated based on the 2^−ΔΔCT^ method (Livak & Schmittgen, 2001). All qPCR primers are listed in Table 6.

### Immune stimulation and 3’ mRNA QuantSeq

FACS sorted RKO-iCas9 or RAW 264.7-iCas9 cells harboring constitutive sgRNA expression (*sgAAVS1*/*sgROSA* or *sgERH*) were treated with 100 or 500 ng/mL dox (Sigma-Aldrich, D9891) for 5 days to induce Cas9-mediated knockouts. Subsequently, they were treated for 4 h with 25 or 10 ng/mL IFNγ (Enzo Life Sciences, ENZ-PRT141-0100 and a gift from G. Adolf, Boehringer Ingelheim, Vienna), 10 or 50 ng/mL IL-1β (Peprotech, 200-01B-10µG and 211-11b), or MQ (non-treated), respectively. Total RNA was isolated as previously described using Trizol, and 3’ mRNA sequencing libraries were generated according to manufacturer’s recommendations (Lexogen, 015-QuantSeq 3′ mRNA-Seq Library Prep Kit FWD for Illumina and 020-PCR Add-on Kit for Illumina). Samples were analyzed on an Illumina HiSeqV4 SR50 platform by the NGS Facility at Vienna BioCenter Core Facilities (VBCF), Austria.

For RNA-seq analysis, quality control was performed with fastqc (v0.11.8) and pre-processing (adapter and quality trimming) with trim_galore (v0.6.2) for RKO and fastp (v0.20.1) for RAW 264.7 cells, respectively. Cell-line specific 3’-ends were determined with 3-GAmES (https://github.com/poojabhat1690/3-GAmES) using human (GRCh38) and mouse (GRCm38) RefSeq and Ensembl annotations for RKO and RAW 264.7 cells, respectively. QuantSeq reads were quantified with SlamDunk v0.3.4 (Neumann *et al*, 2019) using the counting Windows output from 3-GAmES, and 3’-end read counts were collapsed per gene.

Differential expression analysis was done in R (R Core Team (2023)) using DEseq2 (Love *et al*, 2014); genes with log2FC ≥ 1 and padj ≤ 0.05 were considered to have significantly different expression. Drastically different outliers from PCA analysis were excluded from the analysis (RKO samples: replicate 1 – *sgAAVS1* non-treated, replicate 2 – *sgERH* IFNγ treated, and replicate 3 – *sgAAVS1* IL-1β treated). Differentially expressed genes were analyzed for enriched GO terms by Enrichr (Chen *et al*, 2013; Kuleshov *et al*, 2016; Xie *et al*, 2021).

### ERH co-IP and mass spectrometry

Twenty-four hours prior to transfection, 6 x 10^6^ HEK-293T cells were seeded in eight 10-cm dishes (4 negative controls, 4 for ERH-transfection). The cells were transfected with 6 µg plasmid per plate with FuGENE HD (Promega, E2311) as per the manufacturer’s instructions. The other four plates were left untransfected. Media were exchanged 6 h post-transfection. After transfection, cells were grown for 48 hours and harvested by incubation with 1 ml of trypsin/DPBS solution. Trypsin was deactivated by addition of 10 mL of DMEM with FCS and antibiotics, and cells were washed three times with DPBS. Cells lysed by addition of 250 μL 2x lysis buffer (50 mM Tris HCl pH 7.5, 300 mM NaCl, 3.0 mM MgCl2, 2 mM DTT, 0.2 % Triton X-100, 1 tablet cOmplete Mini, Roche) and 150 μL MQ, and by bioruptor sonication (3 cycles, high intensity, 30 sec). GFP-trap beads were prepared as previously described for the Llama antibody against GFP 16 (LaG16) (Fridy *et al*, 2014). The lysate was incubated with the GFP-trap beads for 2 h, washed 5 times with 500 μL wash buffer (25 mM Tris HCl pH 7.5, 150 mM NaCl, 1.5 mM MgCl2, 1 mM DTT, 1 tablet cOmplete Mini, Roche), and boiled at 95°C for 10 min in resuspension buffer (per reaction: 7.5 μL of 4X LDS sample Buffer, 3 μL DTT, 14.5 μL MQ).

Peptides were separated on a 20-cm self-packed column with 75 µm inner diameter filled with ReproSil-Pur 120 C18-AQ (Dr.Maisch GmbH) mounted to an EASY HPLC 1000 (Thermo Fisher) and sprayed online into an Q Exactive Plus mass spectrometer (Thermo Fisher). We used a 94-min gradient from 2 to 40% acetonitrile in 0.1% formic acid at a flow of 225 nl/min. The mass spectrometer was operated with a top 10 MS/MS data-dependent acquisition scheme per MS full scan. Mass spectrometry raw data were searched using the Andromeda search engine (Cox *et al*, 2011) integrated into MaxQuant suite 1.5.2.8 (Cox & Mann, 2008). In both analyses, carbamidomethylation at cysteine was set as fixed modification, while methionine oxidation and protein N-acetylation were considered as variable modifications. Match-between-run option was activated. Prior to bioinformatic analysis, reverse hits, proteins only identified by site, protein groups based on one unique peptide, and known contaminants were removed. For further bioinformatic analysis, the LFQ values were log2-transformed and the median across replicates was calculated. This enrichment was plotted against the – log 10-transformed P values (Welch’s t-test) using the ggplot2 package in the R environment.

### Immunoprecipitation

Cells were lysed in Frackelton buffer (10 mM Tris-HCl pH 7.4, 50 mM NaCl, 30 mM Na_4_P_2_O_7_, 50 mM NaF, 2 mM EDTA, 1% Triton X-100, 1 mM DTT, 0.1 mM PMSF, and 1x protease inhibitor cocktail). Lysates were rotated for 5 min at 4°C and then centrifuged for 10 min at 20,000 x g at 4°C. Supernatant protein concentrations were determined by Pierce BCA Protein Assay Kit (Thermo Fisher Scientific, 23225), and 500 μg total protein was used for each IP (20 μg; 4% was taken for input). IPs were incubated rotating overnight at 4°C with either IgG Isotype control (1:300, Cell Signaling Technology, 2729) or anti-ERH (1:100, Abcam, ab166620). Magnetic beads (Pierce Protein A/G Magnetic Beads, Thermo Fisher Scientific, 88803) were washed 2x in Frackelton buffer, blocked by rotating for 1 h at 4°C in 3% BSA in Frackelton buffer, washed 3x in Frackelton buffer, and 25 µL beads were then added to each IP and left rotating for 2 h at 4°C. After five washes in Frackelton buffer, the IPs were boiled at 95°C for 10 min in Laemmli sample buffer with 10% β-mercaptoethanol, and the whole IP/input was analyzed by WB.

### Immunofluorescence microscopy

2.5 x 10^5^ cells were seeded onto coverslips and fixed with 4% paraformaldehyde (PFA) for 15 min, 48 h post-seeding. Cells were permeabilized with 0.25% Triton X-100 in PBS for 5 min, and blocked in 1% BSA for 30 min at RT. Coverslips were first incubated with anti-ERH antibody (Abcam, ab166620) 1:500 in 1% BSA for 1 h at RT, then with anti-rabbit IgG Alexa Fluor 594 (Invitrogen, A-11012) 1:1000 in 1% BSA for 1 h at RT, and finally with 0.4X Hoechst (Thermo Fisher Scientific, H3569) in PBS for 5 min at RT. Coverslips were mounted using ProLong Gold Antifade Mountant (Invitrogen, P36934) and images were collected using a modified Imager.Z2 microscope (Zeiss) at 63X magnification. Images were modified uniformly across the image using Fiji (Schindelin *et al*, 2012).

### Subcellular fractionation and 3’ mRNA QuantSeq

FACS-sorted RKO-iCas9 cells with constitutive sgRNA expression (*sgAAVS1*, *sgERH*, *sgALYREF*, *sgSRSF1*, *sgMAGOH*, or *sgPOLDIP3*) were treated with 100 ng/mL dox (Sigma-Aldrich, D9891) for 5 days. For subcellular fractionation, 2 x 10^6^ cells were gently lysed in REAP buffer (0.1% NP-40 in 1x PBS) at 4°C, and ≥ 50% of the lysate was taken for the whole cell fraction. Nuclei and cytoplasmic fractions were separated twice through centrifugation at 3,000 x g for 1 min at 4°C. The cellular lysate fractions were analyzed by WB as described previously or used for RNA extraction. RNA isolation was performed with High Performance RNA Isolation kit (Vienna, VBC Molecular Tools Shop, RNAisomag) and a KingFisher Flex robot (Thermo Fisher Scientific). Preparation of cDNA was performed as described above, and qPCRs were run with Luna Universal qPCR Master Mix (NEB, M3003). To assess cryptic splicing, RT-qPCR amplicons were loaded on 2% agarose gels, imaged with a blue-light Safe Imager (Thermo Fisher Scientific), and analyzed with Fiji (Schindelin *et al*, 2012).

Generation of 3’ mRNA sequencing libraries was done according to manufacturer’s recommendations (Lexogen, 015-QuantSeq 3′ mRNA-Seq Library Prep Kit FWD for Illumina, 081-UMI Second Strand Synthesis Module for QuantSeq FWD, and 020-PCR Add-on Kit for Illumina). Samples were analyzed on a NovaSeq SP SR100 platform by the NGS Facility at VBCF, Austria.

For RNA-Seq analysis, reads were pre-processed with fastp (v0.20.1) which was configured to extract 6 nt UMIs and trim 4 nt from the read 5’-ends along standard quality and adapter trimming settings. Quality control was performed with fastqc (v0.11.8). Reads were mapped to 3’-ends with the nf-core/slamseq pipeline v0.1. For the 3’-end annotations we used the previously published set from (Muhar *et al*, 2018) but extended it with GENCODE annotations for missing genes. Read and UMI counts were collapsed per gene and counted with a custom Python script. Differential expression analysis was done in R (R Core Team (2023)) with the RStudio IDE (Posit team (2023)) and by using DEseq2 (Love *et al*, 2014) with apeglm shrinkage option (Zhu *et al*, 2019). Genes with log2FC ≥ 1 and padj ≤ 0.05 were considered significantly differentially expressed. Biomart (v2.50.3) (Durinck *et al*, 2009, 2005) was used to retrieve the number and lengths of exons and introns of canonical transcripts. Hierarchical clustered distance heatmaps and PCA plots were generated with ggplot2 (Wickham, 2016) based on the variance stabilizing transformed (vst) mean mRNA expression of sample groups (Love *et al*, 2014). Enriched GO terms were obtained as described above.

### Poly(A)-enriched RNA-Seq

FACS-sorted RKO-iCas9 cells with constitutive sgRNA expression (*sgAAVS1* or *sgERH*) were treated with 100 ng/mL dox (Sigma-Aldrich, D9891) for 5 days to induce Cas9-mediated knockouts. Cells were treated for 4 h with 25 μg/mL CHX (Sigma-Aldrich, C1988) in DMSO, or DMSO only. Poly(A)-enriched RNA-Seq libraries were generated using the NEBNext Ultra II RNA Library Prep Kit for Illumina (New England Biolabs, E7770), and analyzed on a NovaSeq S4 PE150 XP platform by the NGS Facility at VBCF, Austria.

RNA-Seq analysis was performed using the nf-core/rnaseq pipeline (v3.13.2) (Ewels *et al*, 2020; Harshil Patel *et al*, 2024), which was run on the CLIP Batch Environment (CBE, VBC, Vienna) high performance cluster with Singularity (Kurtzer *et al*, 2017). Briefly, adapter and quality trimming was done with fastp (Chen *et al*, 2018), removal of rRNA reads with SortMeRNA (Kopylova *et al*, 2012), alignment and quantification of reads with STAR (Dobin *et al*, 2013) and salmon (Patro *et al*, 2017), sorting and indexing with SAMtools (Danecek *et al*, 2021), duplicate read marking with Picard, transcript assembly and quantification with StringTie (Kovaka *et al*, 2019), assembly of coverage files with BEDTools (Quinlan & Hall, 2010), and extensive QC with RSeQC (Wang *et al*, 2012), Qualimap (Okonechnikov *et al*, 2016), dupRadar (Sayols *et al*, 2016), preseq (Daley & Smith, 2013), and DESeq2 (Love *et al*, 2014). The genome used for the analysis was built from the latest Ensembl version (GRCh38, v110). Differential gene expression analysis was done as previously described using DEseq2 (Love *et al*, 2014; Xie *et al*, 2021).

IR events were detected and analyzed for differential expression with IRFinder-S (v. 2.0.0) and DEseq2 (Middleton *et al*, 2017; Lorenzi *et al*, 2021) using standard settings. Log2FC (≥ 1) and padj (≤ 0.05) cut-offs were used to determine significance. Specific IR examples were retrieved with the integrative genomics viewer (IGV) (Robinson *et al*, 2011). IR events were considered adjacent if separated by one exon, and a random control was included that selected the same number of introns for each gene as reported previously (Middleton *et al*, 2017). Hierarchical clustered distance heatmaps, enriched GO terms, and canonical transcript information were obtained as described above.

For comparison of intrinsic features, IR events caused by the *ERH* KO were compared to the same number of randomly sampled introns in other genes without intron retention in either knockout (mean IR ratio ≤0.05 in both *sgAAVS1* and *sgERH*) or with intron retention in both knockouts (mean IR ratio ≥ 0.1 and absolute log2FC ≤ 0.2 between knockouts). GC-content was analyzed with the Biostrings package (H. Pagès, 2017). The Maximum Entropy Score was used to calculate 5′ and 3′ splice site strengths using MaxEntScan (Eng *et al*, 2004). Motif enrichment analysis was performed using the MEME suite’s simple enrichment analysis (SEA) (Bailey & Grant, 2021; Bailey *et al*, 2015) and the CISBP RNA motif database (Ray *et al*, 2013). The first 200 bp of the 5’ and 3’ intronic regions (excluding the 6 bp and 30 bp parts of splice sites) and the first 100 bp of the flanking 5’ and 3’ exons (excluding the 3 bp and 1 bp parts of splice sites) were used for the analysis; introns and exons shorter than 200/100 bp were excluded from the analysis. All plotting was done in R using ggplot2 (Wickham, 2016).

## Acknowledgements

All RNA-seq was performed by the Next Generation Sequencing Facility at the Vienna BioCenter Core Facilities (VBCF), member of the Vienna BioCenter (VBC), Austria. Cell sorting for the genetic screens was performed at the IMP BioOptics flow cytometry facility. All other flow cytometry analyses were performed at the BioOptics FACS Facility at the Max Perutz Labs using the Max Perutz Labs instrument pool; we particularly acknowledge Kitti Csalyi, Thomas Sauer, and Johanna Stranner for expert support. Microscopy was performed at the Max Perutz Labs; we thank Irmgard Fischer for her expert support and training. We thank Michael P. Rout for providing a LaG16 nanobody plasmid, and the Protein Production Core Facility (PPCF), IMB, Mainz, Germany for GFP-trap beads preparation. We thank Niels Gehring, Volker Böhm, Sebastian Falk, and Clemens Plaschka for expert advice, discussions, and feedback on the manuscript. We are grateful to the ‘Signaling Mechanisms in Cellular Homeostasis’ doctoral program community, in particular Thomas Decker, Pavel Kovarik, and their lab members. We are grateful to Laura Boccuni for technical expertise and help. We thank Life Science Editors for editing services.

## Funding sources

This work was funded by Stand-Alone grants (P30231-B, P30415-B, P36572, P36945), Special Research Grant (SFB grant F79), and Doctoral School grant (DK grant W1261) from the Austrian Science Fund (FWF) to GAV. MV, M.deA, and SS are the recipients of a DOC fellowship of the Austrian Academy of Sciences. Research at the IMP is supported by Boehringer Ingelheim and the Austrian Research Promotion Agency (Headquarter grant FFG-852936). For open access purposes, the authors have applied a CC-BY public copyright license to any author accepted manuscript version arising from this submission.

## Funding Statement

The funders had no role in study design, data collection and interpretation, or the decision to submit the work for publication.

## Materials, data, code availability statement

The RNA-seq data in this publication have been deposited in NCBI’s Gene Expression Omnibus (Edgar, 2002) and are accessible through GEO Series accession numbers: GSE274069, GSE274070, GSE274071, and GSE274074. All other data generated or analyzed during this study are included in the manuscript and supporting files. Code used in this study is available at https://github.com/ZuberLab/crispr-process-nf, https://github.com/ZuberLab/crispr-mageck-nf, https://github.com/poojabhat1690/3-GAmES.

## Author contributions

Conceptualization: G.A.V., A.S., M.V., J.Z., R.K., F.B., S.A., and M.M-M.; Methodology: A.S., M.V., M.deA., Ni.P., Na.P, R.K., P.A-R., and S.S.; Software, A.S., Ni.P., P.A-R., E.N., and M.deA.; Formal Analysis: A.S., M.V., Ni.P., E.N., and M.deA.; Investigation: A.S., M.V., M.deA., and Na.P., and P.A-R; Resources: M.deA, J.Z.; Data Curation: A.S., Ni.P., and E.N.; Writing – Original Draft, A.S. and G.A.V.; Writing – Review & Editing: A.S., M.V., M.deA., Ni.P., Na.P., P.A-R, S.S., E.N., F.B., R.K., S.A., M.M-M., J.Z., G.A.V; Visualization: A.S.; Supervision: G.A.V.; Project Administration: G.A.V.; Funding Acquisition: G.A.V.

## Declaration of interests

SLA is a co-founder, shareholder, scientific advisor, and member of the board of QUANTRO Therapeutics.

---

**Figure EV 1.**
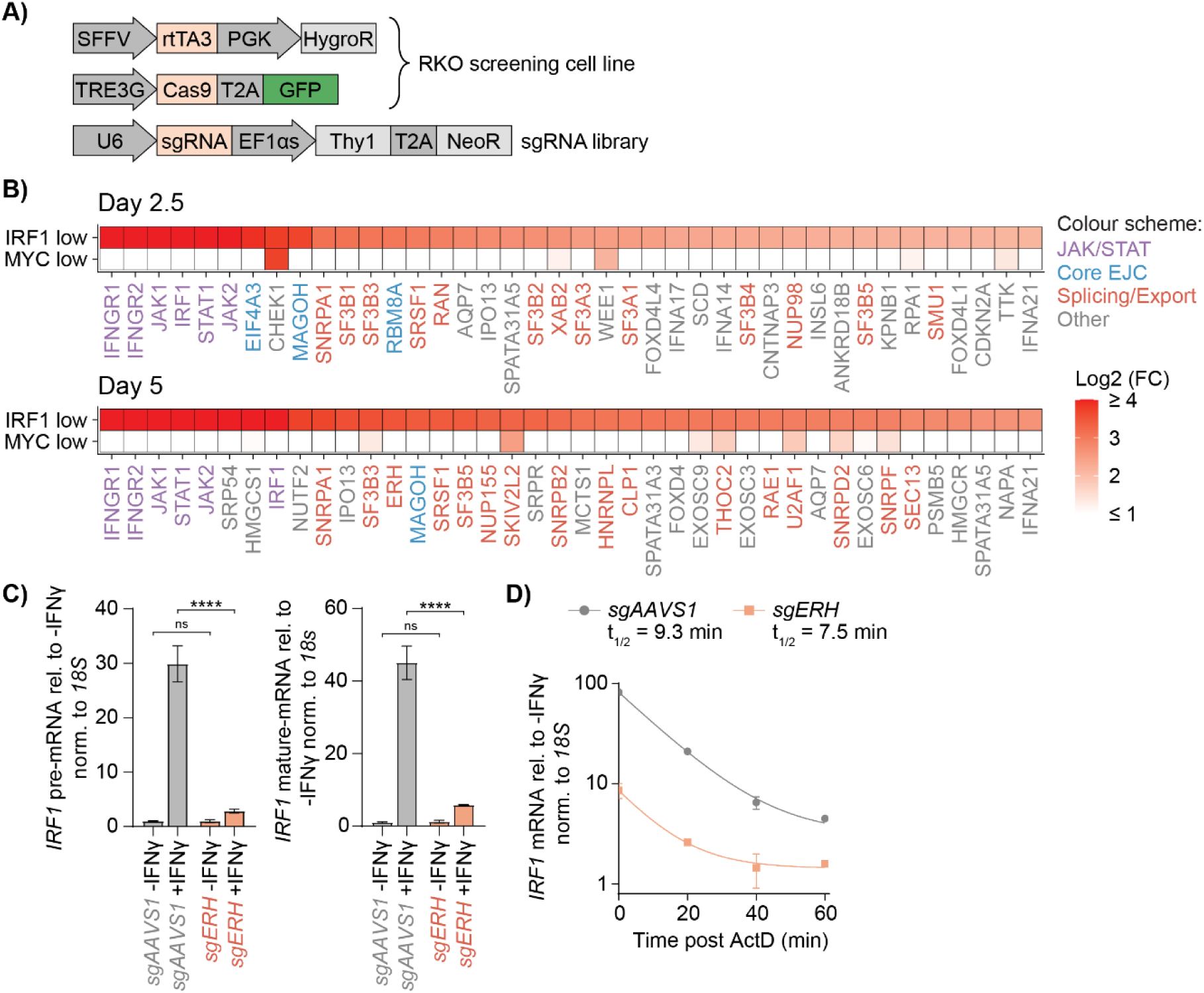
Identification of ERH and other novel positive regulators of IFNγ signaling by genome-wide genetic screening. **(A)** Constructs expressed in the RKO-iCas9 cell line used for the genetic screen. (**B**) Heatmaps showing enrichment of the top 40 positive IRF1 regulators in RKO cells at 2.5 or 5 days of Cas9 induction from the IRF1 genetic screen (Fig. 1D) or a comparable genetic screen for MYC regulators (de Almeida *et al*, 2021). Regulatory factors are color-coded for involvement in: JAK/STAT signaling (purple), EJC (blue), splicing and export (orange), or other processes (gray). (**C**) RKO-iCas9 cells expressing the indicated sgRNAs were induced with dox for 5 days, treated for 4 h with IFNγ, after which pre- and mature *IRF1* mRNA levels were measured by RT-qPCR. Data represent means and sd; n = 3 biological replicates. Two-tailed t-test with Benjamini-Hochberg correction (*p ≤ 0.05; **p ≤ 0.01; ***p ≤ 0.001; ****p ≤ 0.0001). (**D**) RKO-iCas9 cells expressing the indicated sgRNAs were induced with dox for 5 days, stimulated for 4 h with IFNγ, after which actinomycin D was added for the indicated times, and *IRF1* mRNA levels and half-life analyzed by RT-qPCR. Data represent means and s.d.; n = 3 biological replicates.

**Figure EV 2.**
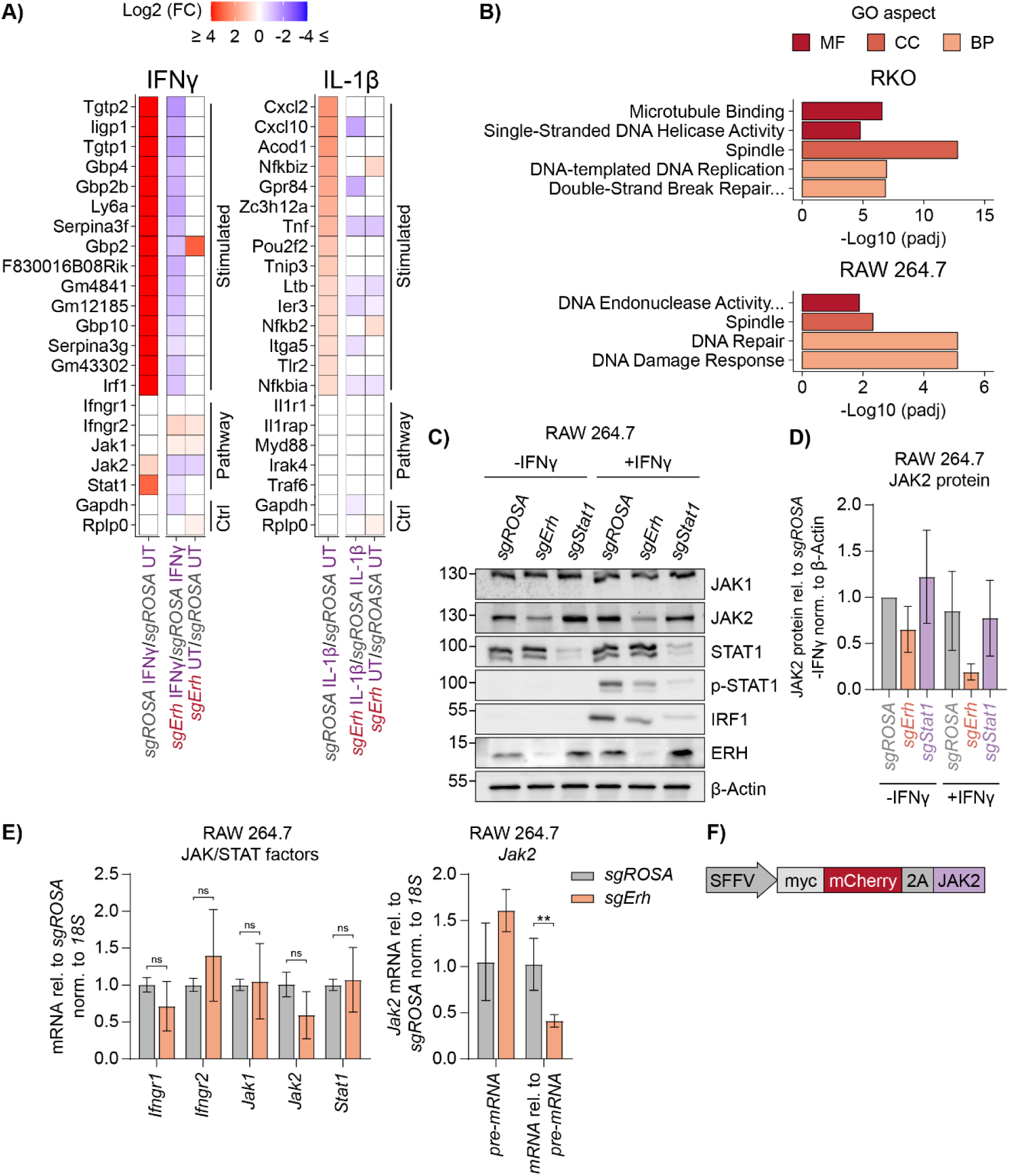
ERH is essential to maintain *JAK2* mRNA and protein levels, and consequently critical for IFNγ signaling. **(A)** Cas9 was induced for 5 days in RAW 264.7-iCas9 cells expressing the indicated sgRNAs, stimulated for 4 h with IFNγ or IL-1β, after which RNA levels were quantified by 3’ QuantSeq. The top 15 stimulated genes and pathway components are shown. Non-significant changes (padj ≤ 0.1) were given a 0-fold change difference. n = 3 biological replicates. (**B**) Selected top enriched GO terms for the subset of mRNAs with reduced expression (log2 fold change ≤ -1 and padj ≤ 0.05) from *ERH* ablation at non-stimulated conditions by 3’ mRNA-Seq in RKO-iCas9 and RAW 264.7-iCas9 cells. MF, molecular function; CC, cellular component; BP, biological process. (**C**) Dox-induced RAW 264.7-iCas9 cells expressing the indicated sgRNAs were analyzed by WB, and (**D**) signal was quantified. Data represent means and sd; n = 5 biological replicates. (**E**) RAW 264.7-iCas9 cells with the indicated genes targeted for 5 days were analyzed by RT-qPCR. Data represent means and sd; n = 3 biological replicates. Two-tailed t-test with Benjamini-Hochberg correction (*p ≤ 0.05; **p ≤ 0.01; ***p ≤ 0.001; ****p ≤ 0.0001). (**F**) Construct used for constitutive *JAK2* cDNA expression in RKO-iCas9 cells.

**Figure EV 3.**
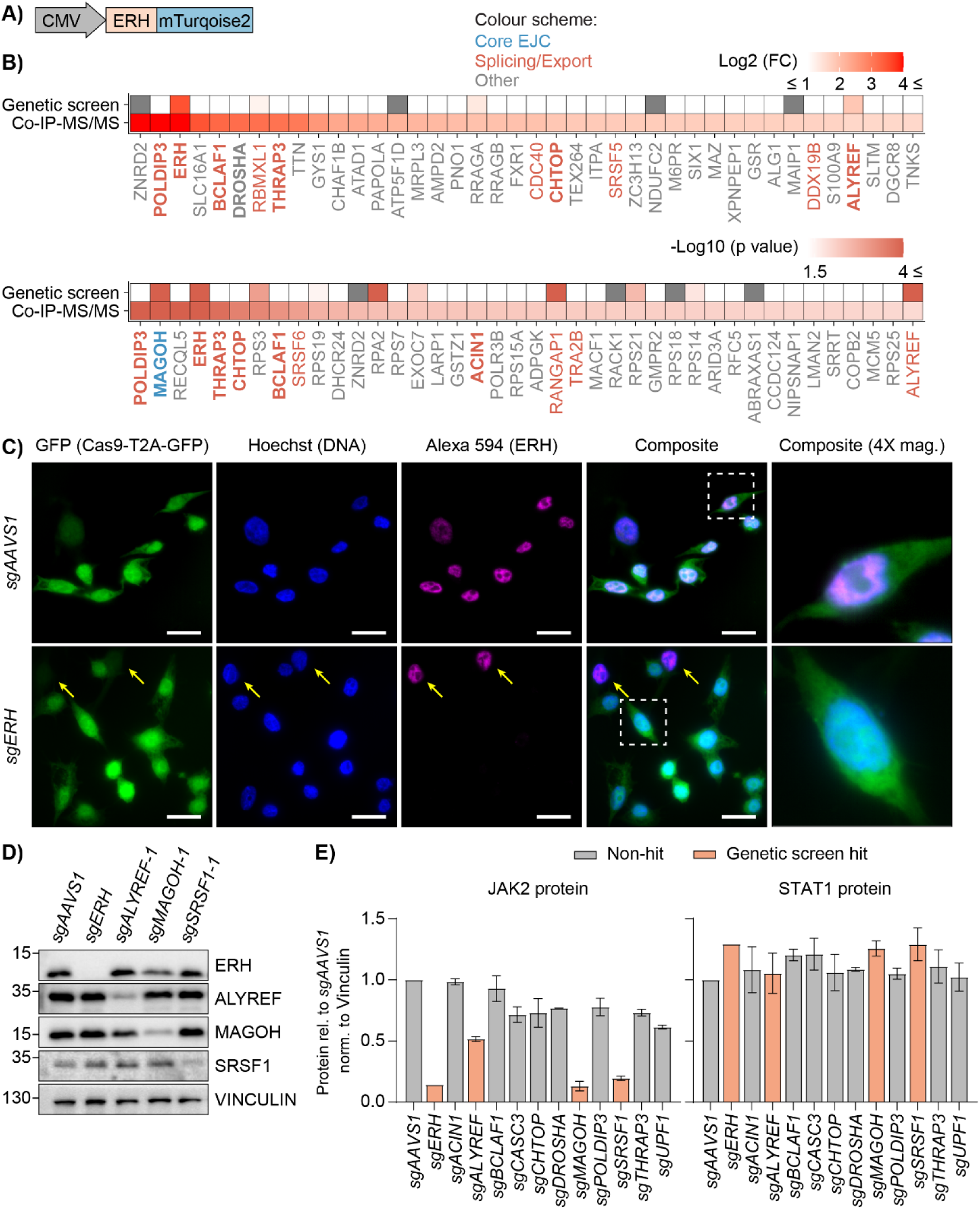
EJC-associated splicing and export factors interact with ERH and are critical for JAK2 production. **(A)** Construct used for ERH co-immunoprecipitation (IP) and tandem mass spectrometry (MS/MS) in HEK-293T cells. (**B**) Heatmaps showing the top 40 enriched proteins in ERH co-IP-MS/MS based on fold change or adjusted p value, and corresponding enrichment in the genetic screen of IFNγ regulators (Fig. 1C). The timepoint (Cas9 induction) with lowest p value or highest fold change from the genetic screen is shown. Factors are color-coded for involvement in: EJC (blue), splicing and export (orange), or other processes (gray). (**C**) Representative immunofluorescent light microscopy images (scale bar, 50 µm) of RKO-iCas9 cells with indicated knockouts (left side) after 5 days of Cas9 induction. Co-expressed GFP was used to mark the cytoplasm and nucleus, whereas DNA (Hoechst) exclusively marked the nucleus. 4X magnification close-up views that visualize the sub-cellular localization of ERH are indicated with dashed white lines. Yellow arrows indicate non-knockout cells in *sgERH* samples. (**D**) Immunoblot from lysates of RKO-iCas9 cells with indicated knockouts after 5 days of Cas9 induction. (**E**) Quantified JAK2 and STAT1 protein levels from (Fig. 3E). Data represent means and sd; n = 2 different sgRNAs.

**Figure EV 4.**
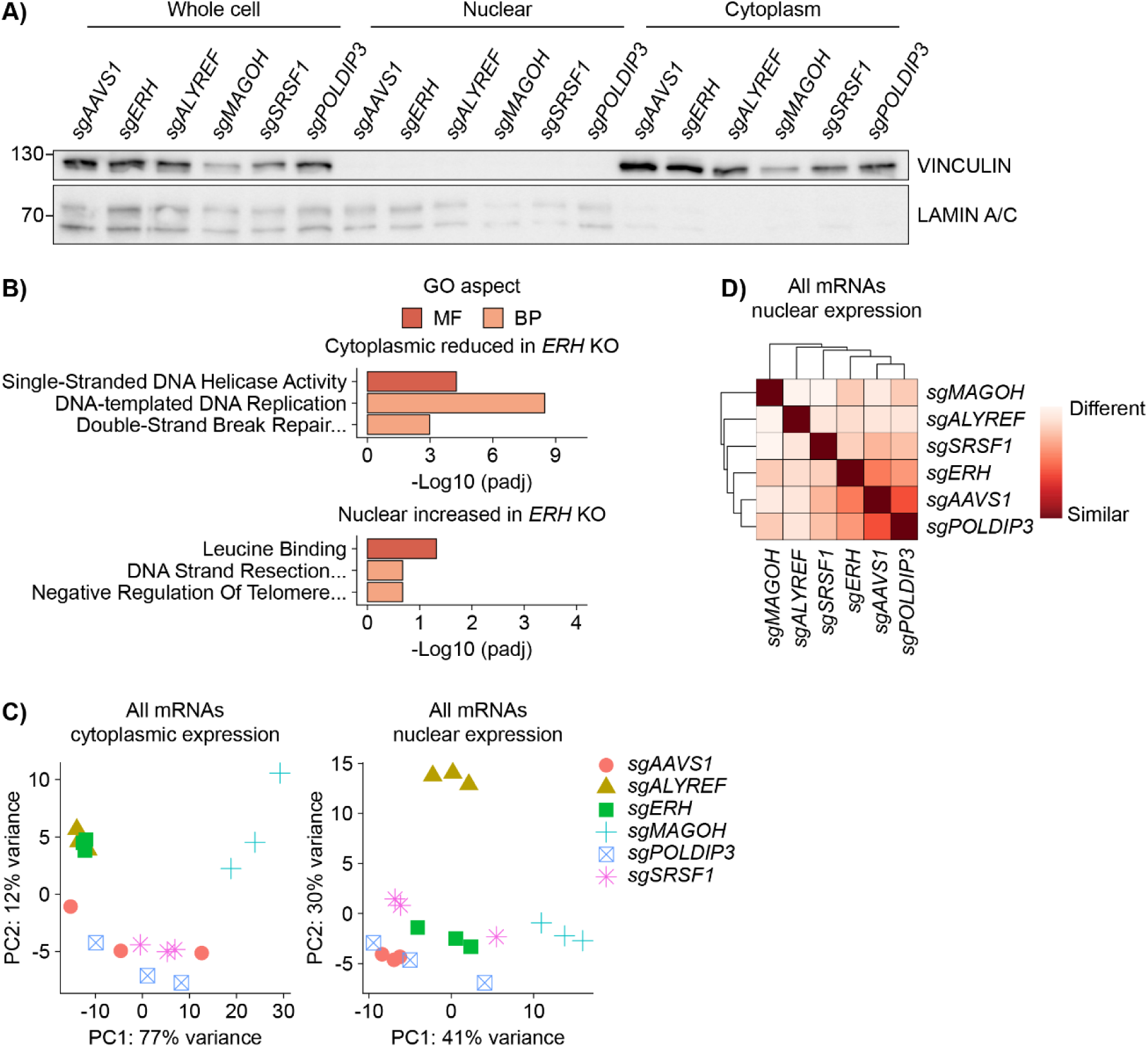
ERH prevents faulty post-transcriptional *JAK2* mRNA processing. **(A)** RKO-iCas9 cells with the indicated targeted genes used for 3’ QuantSeq were fractionated and subsequently analyzed by WB. (**B**) Selected top-enriched GO terms for mRNA subsets (Fig. 4A). MF, molecular function; BP, biological process. (**C**) Principal component (PC) plots of cytoplasmic or nuclear mRNA expression from 3’ QuantSeq samples. (**D**) Hierarchical clustered heatmap from 3’ QuantSeq data assessing similarities between knockouts based on nuclear mRNA expression.

**Figure EV 5.**
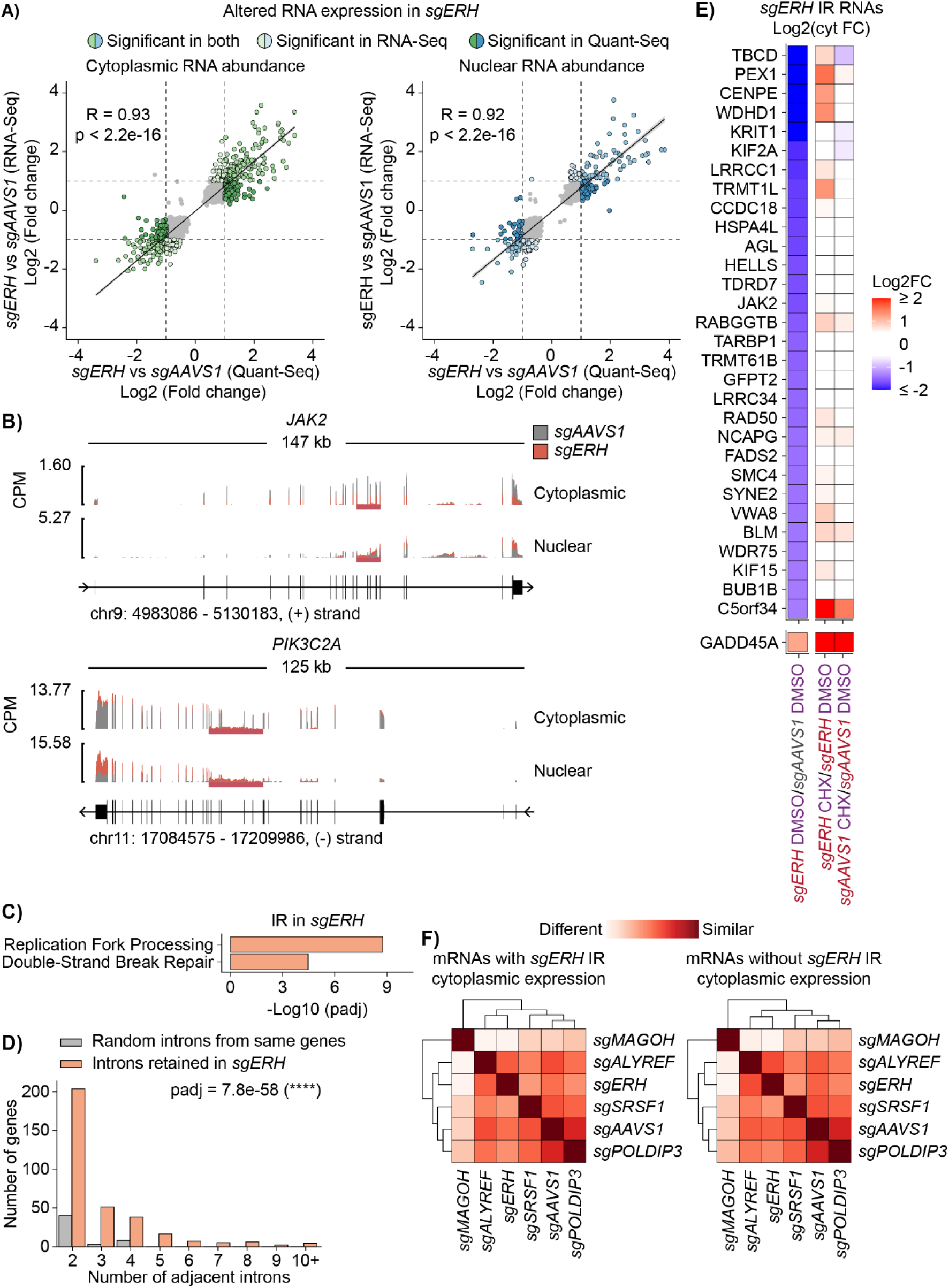
ERH prevents intron retention and disrupted export of *JAK2* and other select mRNAs. RKO-iCas9 cells with the indicated knockouts were fractionated and subjected to poly(A)-enriched RNA-Seq analysis. (**A**) Comparison of altered cytoplasmic (left) or nuclear (right) RNA abundance in RKO-iCas9 cells expressing *sgERH* measured by QuantSeq or poly(A)-enriched RNA-Seq. Only mRNAs with reliably measured differences (padj ≤ 0.01) are shown. Genes with substantial expression differences are highlighted and indicated by dashed lines (absolute log2 fold change ≥ 1). Linear models with sd, and Pearson correlation coefficients and significances are reported. (**B**) *JAK2* and *PIK3C2A* full genes; mean reads per million (CPM) in *sgAAVS1* or *sgERH* are plotted. (**C**) Selected top enriched biological process GO terms for mRNAs with intron retention (IR) in *sgERH* over *sgAAVS1* in RKO-iCas9 cells. (**D**) Correlation analysis of the number of genes containing retained introns that are separated by one exon in *sgERH* samples, compared to randomly sampled introns in the same genes. Two-sided Wilcoxon Rank Sum test (****p ≤ 0.0001). (**E**) Heatmap of subcellularly fractionated poly(A)-enriched RNA-Seq, showing cytoplasmic expression differences for comparisons of RKO-iCas9 cells with indicated knockouts and treatments. The top 30 genes with reduced cytoplasmic expression in *sgERH* that have at least one increased ERH-regulated IR event are shown. (**F**) Hierarchical clustered heatmap depicting the distance between indicated knockout groups based on cytoplasmic expression.

**Figure EV 6.**
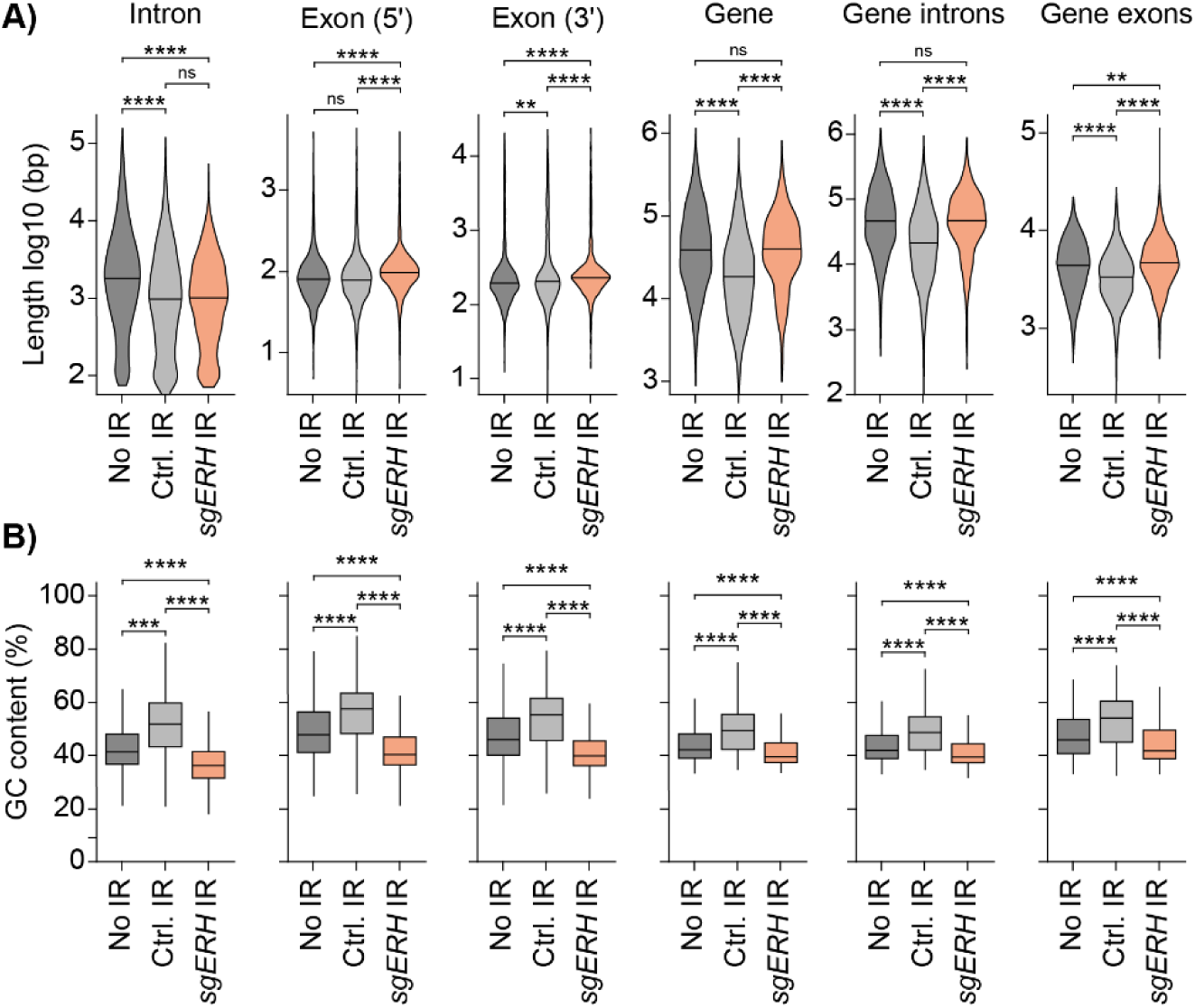
ERH-regulated retained introns and flanking exons are AU-rich. RKO-iCas9 cells with the indicated knockouts were fractionated and subjected to poly(A)-enriched RNA-Seq analysis. (**A**) Violin plots comparing length distributions, and (**B**) box plots comparing GC content, between non-retained (No IR), normally retained (Ctrl. IR), and ERH-regulated retained introns (*sgERH* IR). Genomic regions compared from left to right: introns (Fig. 6A), flanking 5′ exon, flanking 3′ exon, the full gene, all introns in the gene, and all exons in the gene. Two-sided Wilcoxon Rank Sum test with Holm-Bonferroni correction (*p ≤0.05; **p ≤ 0.01; ***p ≤ 0.001; ****p ≤ 0.0001).

## Notes

### Competing Interest Statement

The authors have declared no competing interest.

